# A method to identify high consensus predictions of single-cell metabolic flux

**DOI:** 10.1101/2024.01.15.572211

**Authors:** Michael Amiss, Julian J. Lum, Hosna Jabbari

**Affiliations:** Department of Biomedical Engineering, University of Alberta, Canada; Trev and Joyce Deeley Research Centre, BC Cancer, Victoria, Canada; Department of Biochemistry and Microbiology, University of Victoria, Canada

## Abstract

Altered metabolism is a key contributor to pathology in numerous disease states, including cancer. These changes can occur within certain pathological cells, or within a population of cells. Two recently developed single-cell flux prediction tools, Single-cell Flux Estimation Analysis (“scFEA”) and Compass, have shown success in predicting cellular metabolism using readily available transcriptome data. By adapting the outputs of these tools, we sought to determine if they can work in concert to identify higher confidence consensus flux predictions. We created a set of reaction modules for the two systems. By testing multiple function composites with sets of modularized Compass outputs, we identified a method that showed the highest global similarity to the outputs of scFEA. Our analysis showed broad biological areas of agreement between the results of the two systems when applied to single-cell data arising from both pathological and healthy samples, with pathological samples increasing system consensus. Consensus testing on matched transcriptome and metabolomics data suggested that agreement between the two systems could indicate at least a minimal degree of coherence between both systems and direct metabolite measurements. Overall, we demonstrated that automated Comparisons between the outputs of Compass and scFEA are possible, applicable to data arising from pathological samples, and that such a consensus approach can reveal strongly correlated predictions between these two systems.

**Author summary:** Studying the metabolism of individual cells allows us to understand the mechanisms behind a myriad of diseases. However, single-cell metabolism cannot readily be measured. Computational tools exist to predict metabolism, but validating their outputs requires metabolic measurements. To address this circular shortcoming, we created a method to automatically adapt and compare the outputs of two popular systems used to predict single-cell metabolism from genetic data. In other fields, using predictive methods in an ensemble has provided superior accuracy, and we speculated that the same may hold true for computational predictions of single-cell metabolism. Our work demonstrated that these two systems can be used together to find agreement on a broad range of metabolic processes related to disease. Further, our results, although early, suggest that system agreement may indicate genuine shifts in the underlying biology of a cell population. Owing to the methodologies used by the two systems, such changes could be studied at both a broad or granular level. As our comparison tools provide rapid readouts of such system agreement, they could potentially be used as part of an exploratory pipeline to aid in identification of candidate metabolic mechanisms as drug targets.

## Introduction

In pathological conditions, such as cancer, neurodegenerative disorders, and autoimmunity, metabolism plays a critical role in both disease severity and progression [1–5]. In such pathological states, metabolites are involved in both cell-signaling and proliferation [4].

The study of metabolism at the single-cell level is particularly critical to cancer research. A tumor is comprised of multiple cell types, with varying adaptations to locations and stressors, such as hypoxia [1, 6, 7]. The convergence of these microenvironmental adaptations with the need for increased proliferation results in distinct metabolic profiles amongst tumor cells [1, 6, 7]. Measurements of such metabolic profiles have been reported to provide cancer-specific metabolite signatures [8]. These signatures can potentially identify the originating tumor type of metastatic cells [8]. It is also known that various classes of immune cells within a tumor can undergo extensive metabolic shifts, some of which may contribute to either tumor proliferation or the initial transformation into a malignant state [6].

Once malignancies have been established, tumor cell populations undergo not only separate adaptions to their microenvironments, but to the increased metabolic demands required for uncontrolled growth [1, 9]. In order to maintain a high rate of cell division, malignant cells employ various strategies to obtain large amounts of energy and metabolic precursors required to synthesize vital cell components, including genetic material, lipids and proteins [1, 9]. It is therefore unsurprising that metabolites are a common target of chemotherapeutics [4, 10]. Thus, from both a diagnostic and therapeutic perspective, it is clear that interrogating the metabolism of individual cells may reveal critically important cell-specific characteristics that bulk metabolomics cannot [8, 11].

Despite the importance of such metabolomics data, and the insights it could provide, there is little utility to the data provided by modern single-cell techniques [1, 8, 11]. Currently, single-cell metabolomic measurement protocols suffer from high expense and typically identify only a small fraction, 0.5% or less, of the estimated two-hundred thousand metabolites in the human metabolome [1, 11, 12]. Thus, the advantages of direct single-cell metabolomics cannot yet be realized [1, 11].

In response, several recent methods have been developed to predict the single-cell metabolome from single-cell transcriptomics data [11]. These programs predict metabolic flux, the rate at which metabolites are produced or consumed by a reaction. [13, 14].

Two single-cell flux prediction systems, Single-cell Flux Estimation Analysis (“scFEA”) and Compass, have recently been introduced, and have been used to predict the metabolic states of either tumor or autoimmune cells [1, 5]. These systems have been validated by direct metabolomics, but only in specific domains, and even then, only for highly select processes [1, 5]. However, the markedly different methods used by these tools, as well as their at least partial metabolomics validation, suggests that they could be exploited to their fullest by deploying the tools in concert.

Despite the potential to provide an enhanced level of confidence in single-cell metabolic predictions through a consensus approach, comparing the outputs of scFEA and Compass presents a substantial challenge. The two systems make use of different metabolic models, operate at different levels of reaction resolution, and provide their flux predictions as either flux values or the likelihoods of fluxes being near or at optimum. Further, independent testing or benchmarking of these single-cell flux prediction systems relative to one another appears to be non-existent in any published form. Such testing might suggest the types of originating data or cellular states where the systems agree by way of independent agreement upon with the metabolomics data.

Thus, to address the problem of whether the outputs of the systems can be compared, and the utility of such a comparison, our work attempts to resolve several key questions. The first and most pressing is whether there is a method that will allow for a meaningful comparison between the systems. Second, given that such a method exists, to what extent do the systems find agreement, and at which levels of pathway resolution. Using consensus measures we will attempt to address whether or not there are certain types of data, such originating samples or cells, or pathological states, where these systems agree and on what biological processes. We will also attempt to answer the question of how well these systems agree with each other, and with the metabolomics data, when presented with matched datasets.

## Background

Here, we will introduce the influence of messenger RNA quantity over cellular metabolism and describe how metabolism can be measured. We will discuss how discrete reactions and genes can be related and structured into metabolic models describing the metabolic network within a sample, and how such models are used to estimate metabolism. We will review established methods of estimating the metabolism in a cell or sample. With this, we will discuss two current methods of estimating metabolism at the single-cell level using gene expression inputs, single-cell flux estimation analysis (“scFEA”) and Compass.

### Measuring metabolism

Given the importance of metabolism in the study of diseases, it is frequently interrogated at bulk resolution to determine the chemistry that underlies a pathological state. Metabolites can be directly quantified through technologies such as mass spectrometry (MS) or nuclear magnetic resonance spectroscopy (NMR) [20, 22].

Metabolism can be also be assessed by measuring metabolic flux [5, 20]. Mitochondrial or glycolytic flux can be measured by a Seahorse assay, while labeled isotopic tracing experiments and electron flow assays can provide the necessary data to estimate reaction-level flux [23–26]. However, single-cell metabolic assays are not currently viable due to their limited sensitivity, metabolite coverage and expense [11].

### mRNA and single-cell metabolism

Quantitative RNA-sequencing can be used to count the number of transcripts within the cell as either individual transcript variants or by their common gene of origin [32]. RNA quantities are closely linked to both healthy and pathological cellular states. [32].

Such data provides a path to estimating cell or sample chemistry without the need for metabolomics. The rate of a catalyzed reaction is dependent on the concentration of both the substrate and the enzyme [17–19]. Under the assumption that the substrate concentration is saturating, the concentration of enzyme becomes the rate-limiting reaction component. Thus, as mRNA codes for protein, mRNA levels should, in theory, correlate with the rates of the metabolic reactions that are catalyzed by the enzymes that they code for.

However, bulk RNA-sequencing utilizes samples comprising potentially millions of cells, and this lack of resolution obscures the distinct transcriptomic states of individual cells or cell populations [34]. Recently, single-cell RNA-sequencing (“scRNA-seq”) has been used in the discovery of cell-specific regulatory mechanisms and novel cell types [34], and has been used in the study of heterogenous tumor cell populations and their associated immune populations [32, 35]. Thus, given the maturity of single-cell RNA-seq relative to single-cell metabolomics, and the link between single-cell transcript counts and cell chemistry, single-cell RNA-seq provides a means to broadly estimate the chemistry of an individual cell as either flux or metabolite quantity.

Despite being a more mature technology than single-cell metabolomics, single-cell transcriptome data also presents a challenge. Zero values arising from technical limitations can cause extensive data sparsity, and the stability of a signal arising from mapped single-cell reads is generally reduced relative to bulk [39]. This area has proven to be controversial, as certain zero values may be accurate, and faithfully represent the underlying transcriptomic state of the cell [40]. Measures to impute missing values can include probabilistic methods, which replace missing zeros based on the values in cells with otherwise similar expression, and data smoothing techniques where all values are adjusted based on genetically similar cells [39].

### Metabolic models, flux-balance analysis, and the steady-state assumption

In order to estimate the connected metabolic reactions within an entire cell or sample, metabolic models are required to establish the relationship and connection between genes, metabolites, and reactions [27]. These models are created computationally, drawing on existing databases of metabolic reconstructions. The models can be created using data that includes organism-specific annotations for metabolic genes, experimental data related to metabolic processes, phenotypes, and cellular energetics [27, 28]. Essential to flux prediction at the most basic level, these models can be converted into a stoichiometric matrix that concisely defines the interdependencies between reactions sharing common substrates or products [27, 29–31].

In-silico single-cell flux prediction systems use gene expression inputs, but have their roots in a long-established gene-free method of estimating metabolism: flux-balance analysis (“FBA”). In this method, fluxes can be thought of as variables, with dimensions of moles per unit of dry cellular mass per unit of time, associated with the stoichiometric coefficients of every reactant or product of each reaction or pathway [5, 14].

A stoichiometric matrix is constructed from a metabolic model. The coefficients of each reaction are written column-wise, and the sign of the coefficient corresponds to the production or consumption of a particular metabolite assigned to a row [14]. The matrix is multiplied by a column vector of variables representing fluxes [11, 14]. The resulting rows are linear equations that represent the rate of change of each metabolite (the sum of the fluxes producing or consuming it) which are set to zero under the assumption that the system is at steady state, the condition of unchanging metabolite quantities over time [14]. Linear programming is used to solve for the solution to each equation, while optimizing a desired objective function, such as maximizing cell growth [14]. Each solution then represents the set of fluxes that will produce that optimum [14]. Historically, indirect measures of metabolic activity, such as E.coli growth, or known metabolic properties, have been used to support flux-balance predictions [5, 14].

### Single-cell flux prediction systems

Computational methods used to predict single-cell metabolic flux have only recently advanced [11, 14]. Currently, only a handful of methods applicable to broad metabolic profiling at the single-cell level appear to exist [1, 11]. Optimized solutions to the flux-balance equations can be constrained by single-cell transcriptomics [15]. Such a method, single-cell flux-balance analysis (“scFBA”), has reportedly been used to predict the metabolism and metabolite exchange in the tumor microenvironment of lung and breast cancer cells [15]. In this model, the constraints internal to the stoichiometric matrix, such as flux across cells, as well as upper bounds on each flux variable based on transcript levels are created [14, 15]. However, there is some question as to whether the assumption of a steady-state should be applied to deeply perturbed systems such as cancer cells [1]. Without direct validation, it is unclear whether single-cell flux-balance analysis can accurately model the metabolism of human tumor cells or their associated immune populations.

Recently, two methods, a transcriptome-constrained FBA-based system (“Compass”), and a graph neural network system (“scFEA”) were developed. Both of these systems have had their predictions validated by direct metabolic data. Although such metabolic measurements were limited in scope, and were quantified at the bulk sample level rather than the single-cell level, they suggest that both Compass and scFEA may provide accurate predictions of metabolism at the single-cell level [1, 5, 15].

### Compass

Compass is a recent constraint-based FBA method that uses a penalty matrix derived from the single-cell transcriptome to generate a series of cell-specific objective functions [5]. The system outputs a penalty score for a reaction based on both its calculated flux optimum, and the global expression of metabolic transcripts. Compass does not output a flux prediction, but rather a set of reaction penalty scores where the more penalized reactions are those that are less likely to achieve optimum [5].

The reaction penalty scores are calculated by two algorithms that use the stochiometrically-derived constraints from the *Recon2* metabolic reconstruction. In the first step, the optimal flux for every reaction is estimated by a traditional flux-balance analysis [5]. In this step, the flux through each reaction is set as an objective function to maximize and each resulting linear program is solved. With the optimal fluxes for each reaction calculated, penalty scores are calculated for every reaction in every cell in the second step. A previously created matrix contains a collection of inverse transcript values for which higher transcript results in a lower penalty, and a lower transcript level results in a higher penalty. In the second algorithm, a cell-specific objective function is created by summing every flux variable multiplied by its corresponding penalty matrix coefficient.

Iterating through each reaction, one reaction is held within 95% of the optimum determined by the first algorithm, and the linear program, subject to similar constraints as the first, is solved to minimize the cell-specific objective function (a simplified example is presented in S1 Appendix). The algorithm iterates through each reaction in a cell, and then iterates to the next cell. Conceptually, the reaction penalties generated by the Compass algorithm can be thought of as the minimal global flux profile, subject to transcriptomic effects, required to maintain a high flux through a particular reaction [5]. It is important to note, however, that previously calculated penalties are not used to refine downstream calculations.

Reportedly, Compass was able to predict a large set of altered features in pathogenic Th17 cells relative to healthy controls [5]. Validation of the Compass predictions related to glucose, polyamine and fatty acid metabolism was provided by metabolomic evidence [5]. When assessed against liquid chromatography/mass spectrometry data (LC-MS), the system was successful in identifying a set of increased reactions within both glycolysis and the TCA cycle, which were mirrored in the increased LC-MS abundances for the products of those reactions. The system was also successful in predicting increased beta oxidation during glucose starvation [5].

### Single-cell Flux Estimation Analysis (“scFEA”)

scFEA is an unsupervised graph neural network model that does not rely on either FBA or linear programming methods [1, 36]. The subnetworks of the system use the gene expression associated with their respective reactions to estimate fluxes. The flux outputs of the subnetworks are connected through a factor graph, and a four-term loss function that is minimized during training, primarily by the reduction of flux imbalance across the factor graph [1].

A key distinction between scFEA and Compass is the grouping of related reactions into 171 metabolic modules. The scFEA system uses a metabolic map derived from KEGG pathway data. This metabolic map underwent a network reduction, and was converted to a reduced stochiometric matrix. In the reduction step, clusters of reactions were collapsed into modularized groups of reactions based on their connectedness, and related metabolites were merged. Connected reactions were grouped into a module if the reactions within that module are connected only to each other through a common merged metabolite, or to the incoming substrate or outgoing product metabolite [1]. Each module is further defined as having a single influx and outflux reaction [1, 36]. As a result, scFEA predicts the total flux through a module, and not through its individual reaction components [1].

With the reduced stochiometric matrix, scFEA constructs a factor graph. It is a directed bipartite graph with two types of vertices: variable vertices, which are flux predictions, and factor vertices that calculate the rate of change (the flux imbalance) of a metabolite as a function of the connected variable vertices. The creation and initial use of the factor graph bears some similarity to FBA, as presented our simplified example in “S2 Appendix”. In essence, the reduced stochiometric matrix of scFEA is multiplied by a vector of flux variables, as in FBA. As with FBA, the sum of each row then represents the flux imbalance for one metabolite, as the positively and negatively signed fluxes respectively correspond to metabolite production or consumption. However, with scFEA, instead of setting these sums equal to zero, which would be consistent with the steady-state assumption, each metabolite has an initially unknown flux imbalance.

The flux variables of scFEA are predicted by a collection of subnetworks that take as inputs the metabolic genes that are associated with each module. Each of the factors is represented by the sum of the predicted fluxes that produce or consume its associated metabolite, as mentioned previously. These sums are incorporated into a broader cell and sample-level loss function to be minimized during training. The loss function of scFEA operates to minimize the non-negative summed positive and negative fluxes associated with every metabolite, in other words, minimizing the global flux imbalances or rates of change predicted for all cells in a sample. To achieve this, the system adjusts the weights of each neural subnetwork that estimates a flux, while attempting to enforce coherence between that a particular flux and both specific and global gene expression through additional loss function terms.

While the loss function used by scFEA attempts to approach a steady state condition, it does not reach it. This may be an advantage when studying cancer cell metabolism, since the steady-state assumption may not apply in settings of pathological dysregulation [1]. It is a key distinction between scFEA and flux-balance analysis systems, which universally require the steady-state assumption [1].

One of the strongest pieces of evidence supporting the use of scFEA is its validation by directly measured TCA and glycolysis metabolite abundances. Following a knockdown of a gene known to be involved in DNA damage and hypoxia response in a pancreatic cancer cell line (“Pa03c”), the fold changes in the predicted metabolic fluxes, for the glycolysis and TCA cycle modularized reaction clusters, were calculated for the control and knockdown samples. [1, 37, 38]. The product metabolites associated with these fluxes were measured in the same cell types, and the predicted flux fold changes between the cell types were found to be strongly correlated with the corresponding metabolite fold changes [1].

### The potential for an integrated Compass and scFEA workflow

Beyond the fundamentally different methods of predicting flux, there are several key differences between Compass and scFEA that suggest that their flux predictions may vary when run on similar data.

The consistency between mRNA and flux is critical to the performance of any gene-based flux prediction systems. Both scFEA and Compass use different strategies to make their predictions coherent with mRNA levels at both the reaction and metabolome scale. For Compass, mRNA counts are converted to reaction-specific penalties that are associated with each reaction in a cell-wise objective function. Therefore, mRNA counts affect both the feasible region and the optimal solution for a reaction penalty score. For scFEA, a flux estimate is increasingly penalized as its correlation with both the overall cellular gene expression, and process-level (supermodule) gene expression, decreases.

The gene counts themselves are subject to different pre-processing strategies. Both scFEA and Compass have methods to address the noise and potentially inaccurate zero values that can accompany single-cell data. To counteract this sparsity and technical variation between cells, scFEA can impute missing values by invoking the Markov Affinity-based Graph Imputation of Cells (“MAGIC”) Python package when processing input data. However, this is recommended primarily for 10x data [1]. Compass can employ a k-nearest neighbors (“KNN”) data smoothing method [5].

Given the obvious differences between these two systems, it is uncertain, given their respective metabolomics validation, how they would perform on a common dataset. As mentioned previously, direct measures of metabolite pools, or glycolytic and mitochondrial fluxes, have been used to infer the success of both scFEA and Compass [1, 5]. However, the number of compounds tested was understandably limited, as were the classes of originating cells. Additionally, the precise relationship between instantaneous flux and metabolite concentration may be difficult to determine. While these measures indicate the successes of scFEA and Compass, they have all been at bulk sample-level resolution and have been limited to specific processes.

Thus, it is clear that despite the potential of these systems, there is a deficit in the understanding of their performance outside of their original validation domains, their relative performance when run on common datasets, and whether the two systems can be used as part of a complementary workflow.

### Tandem methods and the potential for increased reliability

Given the methodological differences between Compass and scFEA, and the fact that both systems have been validated by bulk metabolomics, we hypothesized that the two systems could be run in tandem on the same dataset, and the agreement between their flux predictions measured to produce a subset of consensus reactions. We speculated that such reactions may be those for which both systems have accurately predicted a result that could be used for downstream analysis. Here, we developed a method that automatically maps every Compass reaction to every possible scFEA module, and compares the results of both systems across all available biological processes (Figure 1).

**Fig 1.**
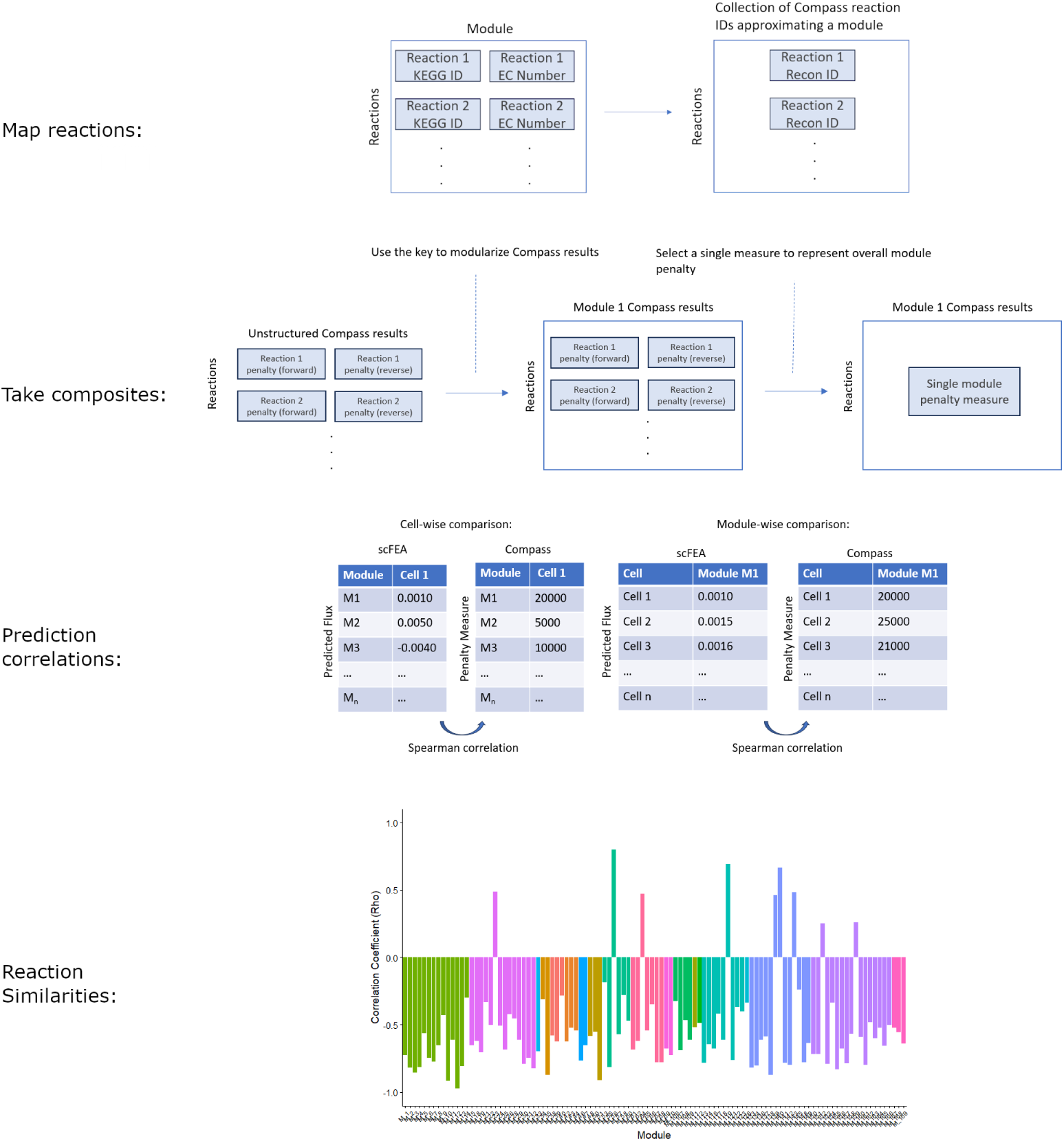
An overview of the consensus methods used in this paper. Compass reactions are mapped to the modularized reaction clusters of scFEA. Once mapped, a composite of Compass outputs (penalties) is taken to represent each module. A correlation is performed between the results for either one module across all cells (and repeated for each module), or all modules across one cell (and repeated for each cell). The module-level results can be visualized to show system agreement, in the form of a negative correlation coefficient, over the entire metabolome.

To the best of our knowledge, no previous attempts at automated comparisons between transcriptome-based flux prediction systems have been published, and large-scale comparisons or benchmarking of single-cell flux prediction systems appear to be similarly absent in literature.

Compass and scFEA predictions operate at different levels of reaction resolution, and the two systems use different metabolic models and reaction identifiers. Given the relatively vast number of directed reaction outputs produced by Compass, it is infeasible to manually map reaction identifiers between the systems. Thus, we automatically mapped Compass reactions into modularized collections approximating those of scFEA. Given that reaction results for each collection spanned a wide range of values, we additionally had to select and test composites of Compass outputs to best represent each module-level flux. As our penalty composites were singular measures derived from multiple reactions, validation at the reaction-level was required to support these findings.

In testing our comparison methods, we attempted to select data and system settings that would both represent typical use-case scenarios which would guide the appropriate application of our methods. To this end, we applied our comparison tests to the Compass and scFEA results produced by bulk and single-cell human cancer and mouse datasets, at both the module and reaction level. To determine the value of consensus reactions relative to experimentally measured metabolomics data, we applied our system to the Compass and scFEA results from two single-cell transcriptomics datasets where matched bulk metabolomics data was available or existed as a published result. This indirectly provided insight into the relative performance of Compass and scFEA on new matched data, as well as suggesting the potential value of consensus reactions.

## Materials and methods

In this section, we describe the methods that we used to determine the similarities between the outputs of Compass and scFEA, as well as the application of these methods to both human and mouse results from these systems. We first detail the command line parameters, versions and libraries that we used when running scFEA and Compass. The next section illustrates how we grouped the reaction identifiers used by Compass into collections approximating those of an scFEA module. In the following sections, we describe the experiments we used to test the performance of eight composite functions that calculate and extract a single Compass measure to represent each module. We further detail our use of the highest performing method to illustrate areas of consensus between the two systems for a broad range of biological processes. The following sections show how we performed additional similarity testing at the reaction-level. In the final sections, we describe how we tested scFEA and Compass on new sets of single-cell data with matched metabolic measurements, and applied consensus measures to the predicted flux results.

### Compass

In this work, Compass version 0.9.10.2 was run under Python 3.8 with CPLEX optimization studio version 20.10 and numpy version 1.21.5. The combination of Python 3.8 and CPLEX optimization studio version 20.10 was chosen because of its documented compatibility with Compass [5]. When Compass was run, the species parameter was set to either human or mouse as appropriate. When k-nearest neighbors smoothing (“KNN”) was enabled, the lambda smoothing parameter was set to 0.25, as in Wagner et al. [5].

### scFEA

Here, scFEA version 1.1.2 was run under Python 3.9 with Pytorch version 1.11.0. The provided human or mouse gene sets and stoichiometric matrices were used as appropriate. Imputation was enabled as necessary during testing, using magic-impute 3.0.0.

### Automating the Comparison of Compass and scFEA results

Because scFEA outputs a single predicted flux through a grouping of connected metabolic reactions (“modules”), and Compass outputs are at the single reaction-level, a comparison between the systems required mapping individual Compass reaction penalties to scFEA modules.

### Re-creation of scFEA reaction module collections with Recon identifiers

Our first goal was to generate a key that could map collections of Recon reaction identifiers, used by Compass, to the reactions within the modules of scFEA. As manually mapping the over 10000 forward and reverse reactions in a typical Compass output to the 171 scFEA human modules was not feasible, we automated the process.

Supplementary table S1 by Alghamdi et al. contains the reactions for 169 modules in their human metabolic map [1]. Metabolic reactions can be identified by multiple naming conventions. This includes the R numbers (“KEGG id”) curated by the Kyoto Encyclopedia of Genes and Genomes (“KEGG”) database and the Enzyme Commission (“EC”) number, which classifies and identifies enzyme-catalyzed reactions [41–43].

Within supplementary table S1 of Alghamdi et al., both the KEGG reaction identifier and EC number for each reaction are listed [1]. To create a minimal set of mappings, we used the KEGG identifier alone. To create the largest set of Recon identifiers possible per module, we used a combined set of both the KEGG identifiers and EC numbers.

Two sets of metadata were required to perform the mappings. These were the reaction metadata (“rxn md.csv”) provided by the Compass GitHub repository and a separate set of metadata from the Virtual Metabolic Human (“VMH”) database containing reaction data for the current version of Recon [5, 44]. The metadata from the VMH database contains both the EC numbers and KEGG identifiers matched to their corresponding Recon reaction identifiers. However, as the EC numbers were not uniquely matched to some Recon reactions in the VMH metadata, we chose to use the Compass metadata exclusively for the EC number and the VMH metadata for the KEGG id. To ensure the broadest possible matches for the KEGG identifiers, we downloaded the data for all of the 13543 reactions available under human metabolism from VMH.

We extracted each collection of modularized KEGG reaction identifiers and EC numbers from the module information in the supplementary table S1 of Alghamdi et al. For each module, we mapped each set of KEGG ids or EC numbers to the corresponding Recon reaction abbreviations within either the Compass or VMH metadata. To create a single list of Recon identifiers for each module, we merged any two sets of module-mappings, originating from either the EC or KEGG identifiers, and eliminated any duplicates. This provided us with a table that listed the Recon identifiers that were uniquely mapped to each of the scFEA modules.

### Comparison of modularized Compass penalties to scFEA predicted fluxes using three datasets

To investigate the similarities between the Compass and scFEA predictions for a broad range of data, we selected three RNA-seq datasets. Because Compass requires gene inputs scaled for library depth, and does not appear to have explicit support for 10x sequencing, we searched for non-10x datasets that were pre-processed with normalization to CPM, TPM or FPKM [5]. As both Compass and scFEA require gene symbol inputs, we additionally chose data with symbol identifiers to avoid potential loss during gene identifier conversion.

Both scFEA and Compass can employ imputation strategies on an input dataset prior to flux prediction [1, 5]. scFEA applies MAGIC imputation directly to the input data, while Compass implements k-nearest neighbors smoothing on the transcript-based penalties it assigns to fluxes. Both of these methods are used to reduce the number of zero values in a dataset, as well as to limit the effects of random noise [1, 5]. Thus, to measure the similarity between Compass and scFEA in samples where the effect of cell-to-cell expression differences were minimized and no imputation was employed, we selected a bulk RNA-seq dataset from an ovarian cancer clonal population (Table 1).

**Table 1.** Three RNA-seq datasets obtained from NCBI geo and processed by Compass and scFEA with consensus and similarity measures applied to the outputs.

| GEO ID | Species | Pathology | Resolution | # of Samples | Technology | Norm. |
| --- | --- | --- | --- | --- | --- | --- |
| GSE123426 | Homo sapiens | Ovarian clear cell carcinoma | Bulk | 12 | Illumina HiSeq 2000 | CPM |
| GSE110499 | Homo sapiens | Multiple myeloma | Single-cell | 174 | Illumina HiSeq 2500 | TPM |
| GSE122330 | Mus musculus | None | Single-cell | 130 | Illumina HiSeq 2500 | TPM |

We obtained two additional datasets to investigate the similarity of Compass and scFEA predictions across single-cell counts. As studying cancer cell metabolism appears to be a major potential application of these systems, we obtained a multiple myeloma dataset for comparison (Table 1). Murine data is broadly used for research purposes, and was the origin of the Th17 sequencing data used Wagner et al. Additionally, the scFEA mouse model appears to use an identical number of modules with similar genes to the human model. Therefore, to test whether the consensus measures could be applied to single-cell mouse data, we chose an additional dataset arising from healthy fetal mouse lung tissue (Table 1). This data also served as a healthy control.

### Post-processing of Compass forward and reverse reaction penalties

Compass outputs both forward and reverse penalties for each reaction, and thus, this posed a challenge when comparing the two systems [5]. To compare the bidirectional reaction penalties produced by Compass to the single predicted fluxes of scFEA, we tested two methods of converting these reaction penalty pairs to a single penalty measure.

The first method we chose was to use the forward penalty (“positive penalty”) only. Since the net movement through a pathway will depend on the forward fluxes of the reactions involved, they are expected to correlate with the net flux through the pathway. As an additional step to explicitly take into account net flux through a reaction, we chose to use ratios of the forward and reverse reaction penalties. The Compass algorithm appears to only produce a penalty of zero if the optimal flux calculated in the transcript-agnostic first FBA step is zero [5]. Therefore, a pseudopenalty of 1 was assigned to such values in the reverse reactions, under the assumption that they would not affect the forward reaction. The penalty ratio itself was calculated as follows:

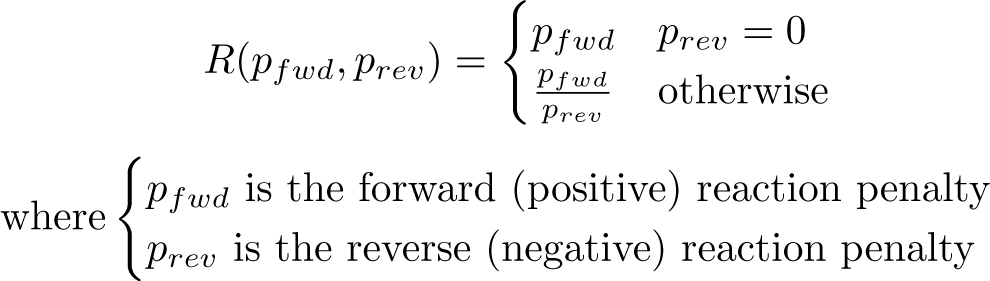

This yielded a ratio for which forward movement through a reaction was expected to be favorable (relative to the reverse reaction) for a lower penalty ratio, and more restricted for a higher penalty ratio. For each of these Compass post-processing methods, we evaluated their success using the downstream similarity measures.

### Penalty measure selection to represent module

To correlate the Compass reaction penalties with the scFEA module fluxes, we first needed to build a table of Compass-derived penalty measures for every module in every cell. To do so, we extracted and calculated the processed Compass penalties for every modularized collection of reactions used by scFEA. Each positive penalty was extracted, or the ratio score calculated, for every modularized collection of reactions where at least one reaction was mapped to a Recon identifier.

A single per-module Compass penalty measure, or penalty measure composite, was required to compare the modularized collections of Compass results to the single module fluxes of scFEA. Using the modularized Compass reaction measures, we obtained either the forward penalty or penalty ratio for each reaction. we took the maximum, minimum or mean value of these measures to test their correlations with the single flux outputs of scFEA. The reactions within an scFEA module are connected, dependent, and must converge to a single outflux [1, 36]. Therefore, the highest Compass penalty measure associated with a module might be expected to most negatively correlate with the scFEA predicted flux through that module. Under the assumption that the reaction-to-module mappings would not be complete, and thus the most penalized reaction could be missed, or that branching within the module could make the previous assumption invalid, the average Compass reaction measure was chosen as an aggregating function as well. The minimum Compass reaction measure was chosen as a control to test the two previous assumptions. The least penalized reaction should be the least inhibitory, and was expected to show far less of a negative correlation with an scFEA predicted flux than those values obtained by the previous two aggregating functions.

With a table of maximum, mean, or minimum penalty measures for every module in every cell, correlations with the scFEA results were possible.

### Cell and module Compass-derived score to flux similarity measure

Alghamdi et al. investigated flux variations both within cells (“cell-wise”) and for specific modules across cells (“module-wise”) [1]. To capture the consensus for flux predictions within and amongst cells, we also used similar cell-wise and module-wise comparisons.

To test the similarity between the results for each cell, a Spearman correlation was performed between the Compass-derived penalty measures for each included module, selected or combined by one of the aggregating functions, and the scFEA predicted fluxes for each included module. We used Fisher’s method to calculate a combined p-value for the results, and Fisher’s Z-transf<u>orm</u> to calculate an average Spearman’s *ρ* by applying the following formula: 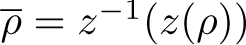 (where *z*(*x*) is Fisher’s Z-transform).

To determine which modules showed the strongest significant correlations between the Compass and scFEA results, we performed a Spearman correlation between each combination of aggregating function and module penalty results derived from Compass, and the scFEA predicted flux, for each module across all cells. To identify the significantly correlated modules, we took the subset that had a p-value of less than 0.05. Modules with a static scFEA predicted flux, or Compass penalty measure, for which no correlation was possible were counted as NA. Because penalty measures were expected to negatively correlate with the predicted net flux through a module, the subset of modules producing significant, but positive, correlations were counted as an additional measure of Compass-to-scFEA dissimilarity. The ratio of positive to significantly correlated modules was taken to measure overall module-wise consensus.

Each of the 22 supermodule classes of scFEA is associated with a broad description of the biological processes it encompasses [1]. To investigate the biological function of the modules for which there was a significant Compass to scFEA correlation, we extracted their corresponding supermodule identifiers from the supplementary table S1 of Alghamdi et al. [1]. To attach a biological process to each module in the correlation results, we created table of supermodules ID, their description, and their associated modules based on the supplementary tables and figures of Alghamdi et al. [1]. The modules in each set of correlation results were assigned a supermodule ID and description based on the supermodule table.

### Compass reactions to scFEA module flux correlations

To investigate the similarities between specific Compass reaction penalties and scFEA predicted fluxes, and to investigate whether the limited Recon to KEGG id or EC mappings reduced the accuracy of the maximum positive penalty method, we performed direct reaction to flux correlations as an alternative to cell-wise or module-wise correlations.

Recon2 subsystems are analogous to an scFEA supermodule class in that they provide a broad biological classification of the process that a reaction belongs to. We selected three subsystems, the TCA cycle and glycolysis, both key processes in cellular energy production, and purine synthesis, for a reaction-level investigation. TCA and glycolysis were selected based on their metabolomics validation in both the work of Wagner et al. and Alghamdi et al., as well as the performance in the module-wise comparison here [1, 5]. The purine synthesis subsystem was selected for investigation based on the number of positively correlating modules, within the purine synthesis supermodule class, in the module-level analysis. We calculated the Spearman correlation between each scFEA predicted flux within a supermodule and every forward Compass reaction within its Recon2 subsystem analog. To retain only significantly correlated reactions, those with an associated p-value of less than 0.05 were dropped. For each subsystem, the significant module-to-reaction correlations were extracted from the total correlation matrix.

### Flux prediction and consensus measures for matched RNA-seq and metabolomics data

Both Wagner et al. and Alghamdi et al. validated their flux predictions against bulk metabolomics data [1, 5]. Given the apparent scarcity of matched single-cell gene expression and matched bulk or single cell metabolomics datasets, we decided to test each system on the matched metabolomics and transcriptomics data of the opposing system.

### Compass Th17 data analyzed by scFEA

Using Compass, Wagner et al. predicted upregulated glycolyis, polyamine, and TCA cycle reactions in pathogenic Th17 cells relative to non-pathogenic controls [5]. In validating these predictions, the authors provided their LC/MS metabolomics results from pathogenic and non-pathogenic *Mus musculus* Th17 cells [5]. Although the underlying metabolomics data was not provided, we tested whether the scFEA predicted fluxes for the two populations were qualitatively consistent with the glycolysis, TCA and polyamine metabolomics results of Wagner et al. To determine the agreement between scFEA and Compass for the Th17 data, and to determine their consistency with the metabolomics of Wagner et al., we ran Compass on the Th17 data and applied both the cell-wise and module-wise consensus measures, with the maximum positive penalty method, as described in sections “Automating the Comparison of Compass and scFEA results” and “Comparison of modularized Compass penalties to scFEA predicted fluxes using three datasets”.

### Data Sources and Pre-Processing

The sequencing data analyzed by Wagner et al. arose from 139 Th17 pathogenic cells (“Th17p”) and 151 non-pathogenic (“Th17n”) cells [5]. We obtained the data for these cells at the provided GEO accession (GSE74833). We downloaded the data for the 139 pathological (Il-1β+IL-6+IL-23, sorted for IL-17A/GFP+) cells under the sub-series heading GSE75109 and the 151 non-pathological (TGFβ1 IL6-48h-IL-17A/GFP+) cells under the heading sub-series heading GSE75111.

As these collections contained the gene expression data for each cell in individual files, we joined the FPKM normalized counts for each cell within the sub-series into a single matrix. For compatibility with scFEA and Compass, gene symbols were chosen from the multiple gene identifiers in the data. To avoid information sharing between the pathological and normal Th17 cells when imputation was enabled, the expression data was saved as two different sets.

### scFEA module selection

To compare the scFEA flux predictions to the Compass metabolomics results, we needed to select a set of modules associated with either the pathway in general or the production of each metabolite. Using the module information in the supplementary data table S1 provided by Alghadmi et al., we selected fourteen modules associated with glycolysis and the TCA cycle by their supermodule description [1]. In addition to the differences in these pathways between the Th17n and Th17p cells, Wagner et al. also validated their predictions of disordered polyamine metabolism with metabolomics measurements [5]. To investigate whether scFEA could produce fluxes consistent with the polyamine metabolite figures of Wagner et al., we selected four modules producing (“Compound out”) either putrescine, spermine, ornithine or creatine from the scFEA module information in supplementary table S1 [1, 5].

### scFEA processing and analysis

To obtain flux predictions for both the Th17n and Th17p sequencing data, we ran scFEA with both the Th17n and Th17p count matrices using the included mouse gene model and stochiometric matrix.

For comparison with the Compass polyamine, TCA and glycolysis metabolite figures, we extracted the flux predictions for the 14 glycolysis and TCA modules, and 4 polyamine modules, from the results for either the normal or pathological samples. The significance of the mean differences in fluxes between the two cell types was determined by a Bonferroni-corrected Welch’s t-test. The median and mean differences between the pathological and normal samples were calculated for comparison.

### Compass processing and analysis

For comparison to our scFEA results, we ran Compass on both datasets with KNN smoothing enabled or disabled. Compass was run with four combinations of datasets and parameters (KNN with Th17n, KNN with Th17p and both Th17 and Th17p without imputation). The post-processing code provided by Wagner et al. (“Demo.ipynb”) was adapted for the separate sets of penalties, and used in our downstream analysis [5].

### scFEA and Compass similarity measures

To determine the consensus between the predictions for Compass and scFEA, and to relate the consensus measures to metabolomics measurements, we applied the cell-wise and module-wise consensus measures described in the section “Cell and module Compass-derived score to flux similarity measure” to the pathological and normal Th17 results from both systems.

### Compass analysis of scFEA pancreatic adenocarcinoma dataset

Alghamdi et al. tested their predictions in pancreatic adenocarcinoma (“Pa03c”) cells. Single-cell data from two groups of cells were used, a control group and a apurinic endonuclease (“APEX1-KD”) gene knockdown group. The authors validated their system by calculating the fold changes in the bulk measurements of TCA and glycolysis metabolites between the cell types, and correlating these with the fold changes in the predicted fluxes that produce their respective metabolites [1]. To test the performance of Compass on this matched dataset, we approximated the analysis by Alghamdi et al. by comparing the fold changes in the predicted activity for a reaction producing a metabolite to the fold changes in metabolite levels. To compare the results of the two systems, we approximated the analysis of Alghamdi et al. on the same Pa03c data with scFEA, but used the whole unfiltered dataset and the default hyperparameters. We applied the maximum positive penalty consensus measure described in sections. “Automating the Comparison of Compass and scFEA results” and “Comparison of modularized Compass penalties to scFEA predicted fluxes using three datasets”.

### Data sources and pre-processing

To create subsets of knockdown and control counts, we downloaded the series matrix for the Pa03c normoxic samples from the GEO accession GSE99305. We used the sample characteristics in the series matrix to identify the APEX1-KD cells as “transfection: APE1 siRNA” and the control cells as “transfection: scrambled siRNA”. For each, we created a set of sample identifiers to subset the total counts.

To create the appropriate sample subsets to use as Compass input, we downloaded the processed count matrix for all of the cells. The data was cleaned to remove sequencing metadata from the gene identifier rows. As Compass required a gene symbol identifier and a count matrix scaled for library depth, the AnnotationDbi (1.56.2) and org.Hs.en.db (version 3.14.0) were used to convert the Ensembl identifiers to symbol, and edgeR (3.36.0) was used to calculate the counts per million (“CPM”) for the count matrix. For duplicate genes produced by one-to-many identifier mappings, the CPM counts were summed to a single value. Using the APEX-1 KD or control sample identifiers that we obtained from the series matrix, we created two subsets of the total counts corresponding to these APEX-1 KD or control identifiers.

To compare our Compass outputs to directly-measured metabolite quantities, we obtained the corresponding Pa03c metabolite profiles from the supplementary materials of Alghamdi et al. [1].

### Compass penalty predictions and post-processing

For both the APEX1-KD and control counts, we ran Compass to obtain the calculated reaction penalties. Compass was run with the same version and environment as for the Th17 analysis. We used the settings provided for human scRNA-seq data in the Compass tutorial for processing both datasets [5]. To positively correlate with the metabolites, the resulting penalty matrices were converted to reaction activity scores by taking the inverse penalty with a pseudopenalty of one, as suggested in the post-processing code of Wagner et al. [5].

### Mapping compass reactions to Pa03c metabolites

A comparison between the Pa03c metabolomics data and the Compass output required that we first match the TCA/Citric acid metabolites to a reaction. As the Pa03c metabolites were named using their KEGG compound identifier, we converted the 22 metabolites into their common names using the Metaboanalyst conversion tool [45]. For the two KEGG identifiers (C00031 and C00668) that did not produce a match, we obtained the compound names directly from the KEGG database [43]. From the resulting list of compound names, we identified seven of the eight metabolites used in the predicted flux to metabolite correlation by Alghamdi et al., with the exception of oxaloacetic acid.

To find the forward Recon reaction identifiers associated with the production of each of the seven metabolites, we referenced the canonical TCA and glycolysis pathways (those with established and well-known descriptions) and identified the enzymes associated with the production of the metabolite. To find the associated Recon reaction, we searched for each of the enzymes in the human metabolism section of the Virtual Metabolic Human (VMH) database [44]. From the list of reactions returned, we selected those that matched the canonical forward reaction for that step. For reactions with both cytoplasmic and mitochondrial variants, the compartment was selected based on whether the canonical reaction was associated with glycolysis (cytoplasmic) or the TCA cycle (mitochondrial).

To extend the analysis, we added four other TCA or glycolysis metabolites present in the metabolomics data from Alghamdi et al. in a similar fashion. As sn-glycerol-1-phosphate dehydrogenase did not return any reactions in the VMH database, we used the University of California, San Diego’s Biochemical, Genetic and Genomic (“BiGG”) model database and the Compass reaction key to identify the reaction converting DHAP to Glycerol-3-phosphate [46].

### scFEA predicted fluxes

To compare the scFEA predictions for the Pa03c cells to those of Compass, scFEA was run on both the APEX1-KD and control counts. The default human settings, including the human module genes and stoichiometric matrix were selected as parameters.

### Metabolite to activity and flux similarity

As the bulk metabolomics data consisted of a single per metabolite quantity, the mean activity or predicted flux for each reaction cell was taken for both the APEX-KD and control Compass outputs. Following the analysis of Alghamdi et al., we calculated the fold difference in the mean flux or activity between the control and APEX1-KD cells, as well as the fold difference in the mean product metabolite levels. We measured the similarity between the predicted flux or activity fold changes and their associated metabolites by a Pearson correlation, and plotted the results with the inclusion of a linear model.

## Results

In this section, we will demonstrate how the individual reaction penalties used by Compass can be optimally grouped into modularized collections approximating those used by scFEA. For the three datasets described in section “Comparison of modularized Compass penalties to scFEA predicted fluxes using three datasets”, we will present the results of correlating a single Compass penalty measure with the predicted flux of scFEA for each module. Using these results, we show how the maximum positive penalty method can be used to illustrate areas of biological agreement between the two systems. As an alternate method that does not rely on a single Compass penalty to represent a module, we will demonstrate the results of a reaction-level analysis and show how individual Compass penalties correlate with the flux predictions of scFEA. Finally, we will show the results of testing the performance of each system, and agreement between the systems, when they are run on the matched gene expression and metabolomics data of the opposing system.

### Consensus Measures and Testing

As mentioned in methods section “Compass”, Compass outputs reaction penalty scores for both the forward direction reverse directions, if a reaction is reversible [5]. The predicted flux outputs of scFEA are for a module, an artificial grouping of related reactions with singular and dependent input and output fluxes [1].

### Compass to scFEA Module to key generation

As a first step to determine the extent to which Compass reactions could be associated with an scFEA module, collections of Recon reaction codes corresponding to an scFEA module were generated. The KEGG reaction identifiers and EC numbers corresponding to the reactions within each module were tested separately for their potential to produce unique mappings to a Recon reaction identifier.

The EC number produced more mappings to a Recon reaction code than the KEGG identifiers, but resulted in a greater number of modules where the mapped reactions exceeded the number in the original. The EC number produced 43 modules with zero reaction mappings in contrast to the 52 modules with zero mappings produced by the KEGG-to-Recon mappings. 119 of the reaction collections produced using the EC number contained more than half the number of reactions in the original scFEA module. However, the EC number reaction collections produced 80 modules where the number of reactions exceeded those of the original scFEA reaction set, in contrast to the 22 such reaction collections produced using the KEGG identifier.

A range of modules from 71 to 105 showed zero mappings to a Recon code for either the KEGG identifiers or the EC numbers. The KEGG identifiers and EC numbers that were programmatically obtained from the supplementary scFEA module information for this range were manually investigated. This revealed no valid EC numbers or KEGG identifiers for any module in this range, and such identifiers were solely used as placeholder values. A further investigation of the scFEA module information revealed that modules 71-105 were used exclusively for single transport influxes and are not expected to be associated with any reaction identifier.

Combining the KEGG identifiers and EC number mappings resulted in a higher number of unique Recon reaction identifiers per module. The number of modules with zero mappings was reduced to 37, of which only two modules, 120 and 125, were not in the 71-105 range. Discounting this range, approximately 99 percent of modules were mapped to at least one Recon identifier. For the modules outside of the 71-105 range, 75 percent of modules contained at least half the number of reactions as in the original scFEA module.

To account for the cases where the mapped Recon identifiers exceeded the reactions in the original scFEA module, the reactions were investigated. Although an exhaustive search was not performed, several Recon compartmental reaction variants, such as cytosolic, mitochondrial or peroxisomal reactions, were found to have been mapped to by the same KEGG identifier or EC number. Additionally, reaction variants substituting ITP for ATP were discovered within certain modules, although not to the exclusion of ATP utilizing reactions. Other reaction variants substituting multiple related substrates, such as epimers, were consistent with the reactions in the original scFEA module.

A non-exhaustive manual comparison of four modularized collections of Recon identifiers to the scFEA metadata was performed to determine whether the reaction groupings had been correctly assigned. The reactions within four modules, 1, 30, 40 and 150, and the Recon2 subsystems associated with each reaction, namely glycolysis/gluconeogenesis, methionine and cysteine metabolism, aspartate metabolism and pyrimidine metabolism, were either consistent with the reactant-to-product conversion of the associated scFEA module, or the reaction could be directly located in the original scFEA module information when converted back to a KEGG id.

Collectively, these results support the hypothesis that Recon2 reaction identifiers can be broadly clustered into groups approximating the reaction collections within an scFEA module. This can be achieved through automated KEGG identifier or EC number to Recon identifier mapping for all modules that are not purely related to transport fluxes. The most mappings can be obtained by combining the unique results from both the KEGG identifiers and EC numbers. However, regardless of the method chosen, the mappings may include reaction variants that are not consistent with the compartment or substrates of the original scFEA module.

### Module-to-Reaction Compass to scFEA Comparisons

The previously generated Compass-to-scFEA module key allowed the Compass reaction penalty results arising from the three datasets described in section “Comparison of modularized Compass penalties to scFEA predicted fluxes using three datasets” to be grouped into modules approximating those of scFEA. From these collections, the positive-to-negative penalty ratios were calculated for each set of bidirectional reactions, or the positive penalties alone were taken as a measure of bidirectional reaction likelihood. To test the hypothesis that a consensus could be reached between the results of scFEA and Compass through a single penalty score, or single composite of penalty scores, a per-module measure of Compass penalty was calculated by selecting either a reaction penalty ratio or positive reaction penalty using one of four aggregating functions. This allowed for a one-to-one comparison to each of the predicted scFEA fluxes.

An increasing Compass penalty score implies that optimal flux through a reaction is less likely to occur [5]. Thus, a Compass penalty within a collection of reactions, known to be part of an scFEA module, is expected to negatively correlate with the scFEA predicted net flux through that module if the systems are in agreement on the flux potential of that reaction. Each combination of penalty measure and aggregating function was tested for its ability to produce a negative correlation between the Compass and scFEA results for a particular cell or sample (“cell-wise”). The three datasets described in section “Comparison of modularized Compass penalties to scFEA predicted fluxes using three datasets” were used as input for scFEA and Compass. To test the effects of imputation on the prediction consensus for Compass and scFEA, we selected the multiple myeloma dataset, as it produced the highest module-wise consensus performance in the analyses described in the following section.

### Sample-wise similarity testing

The similarities between one of the Compass penalty measures for each module in a cell or sample, aggregated by one of the four methods, and the predicted scFEA flux for the analogous modules in the same cell or sample, were determined. Across the results for each dataset, weak (*−*0.26 *≤ r ≤* 0.036) cell-wise correlations were observed for any combination of the maximum, sum or mean of positive penalty or penalty ratio (Tables S1-S4). For the bulk clear-cell carcinoma dataset, the lowest significant combined p-values, and the most negative correlation coefficients, were observed using the sum of the penalty ratios in each module (*r* = *−*0.26, *p_comb_* = 4.75 *×* 10*^−^*^19^) (Table S4). For the single-cell results, the maximum positive module penalty demonstrated weak but significant cell-wise correlations in the mouse fetal lung dataset (r=*−*0.084, *p_comb_*=0.0053) and near significant correlations in the multiple myeloma dataset without imputation (r=*−*0.066, *p_comb_*=0.091) (Tables S1, S2). Collectively, this provided support for the idea that a sample-wise consensus between the results of the two systems can best be established for bulk samples by the sum of the penalty ratios. For single-cell data, the results provide weak support for the use of the maximum positive penalty to achieve the highest cell-wise consensus.

### Module-wise similarity testing

The trends in single fluxes across all cells or samples can be compared between Compass and scFEA. Such results could suggest whether or not the systems are capturing the same variation in a particular flux across all of the cells in a sample, or across all of the samples in a set. For the single-cell results, either the maximum, sum or mean of the positive penalties produced the highest numbers of significantly correlated modules (Tables S1-S3). The positive penalty methods showed nearly identical numbers of significantly correlated modules for the multiple myeloma dataset, with or without Compass KNN imputation, and ranged between 82-90 such modules for the mouse lung dataset. Relative to the multiple myeloma results, reduced numbers of significantly correlated modules were observed generally for either the ratio method, or the positive penalty method, in the bulk dataset and the single-cell mouse lung dataset (Tables S2, S4).

Positive correlations are not expected between a Compass reaction penalty and an scFEA predicted flux within the same module. Thus, positive correlation coefficients were used as a measure of disagreement between the Compass derived module penalty measure and the scFEA flux predictions. For the single-cell results, the maximum positive penalty method demonstrated a lower average positive-to-significant module ratio for any non-minimum aggregating function (Tables S1-S3). The lower average was the result of a reduced positive-to-significant module ratio in the mouse dataset (Table S2). In the bulk results, the lowest such positive-to-significant ratios for a non-minimum aggregating function were observed for the sum and mean of the positive penalty (Table S4). However, these were associated with only 23 significantly correlated modules.

When imputation methods were used for both scFEA and Compass on the multiple myeloma dataset, the overall number of significantly correlated modules remained similar. However, the number of significantly positive correlations nearly tripled relative to the non-imputed results (Tables S1, S3).

Thus, given the module results for these single-cell data, but not bulk data, the maximum positive penalty method produced a relatively high, or nearly identical, number of significantly correlated modules and the lowest average percentage of false positives. The results further suggest that enabling imputation in the two systems decreases the consensus, as measured by an unexpectedly positive correlation.

### Module-wise consensus across biological processes

Given the superior performance of the maximum positive penalty method in the consensus testing, it was used to study the module-wise consensus across each dataset or imputation setting by matching the correlations with their biological processes. This was achieved by mapping each module to its supermodule description. To determine the patterns of agreement and disagreement between the two systems, the maximum positive penalty to predicted flux correlations were plotted for each module in each set of Compass and scFEA results. The single-cell results demonstrated significant negative correlations for at least twelve of the fourteen modules in the glycolysis and TCA supermodule class (Figure 2A, 2C). A majority of the pyrimidine synthesis (seventeen total), serine metabolism (eighteen total), urea cycle (eight total) and both fatty acid synthesis modules, were significantly and negatively correlated for both the mouse lung and non-imputed multiple myeloma results (Figure 2A, 2C). At least seventeen of the twenty-two supermodules were represented for each single-cell result, although the sialic acid and glycan synthesis classes were notably absent. In the bulk results, glycolysis and TCA were notably reduced down to a single negatively correlated module (Figure 2D). The transporter modules, 71-105, which could not be mapped to a Compass reaction, were not present in any of the results. Of the significantly correlated modules for the non-imputed multiple myeloma results and the bulk clear-cell carcinoma dataset, the majority had a negative Spearman’s *ρ* with a magnitude of greater than 0.5 (Figures 2A, 2D). However, the bulk data demonstrated a markedly reduced number of significant correlations. The multiple myeloma results that used imputation showed a broad distribution of unexpectedly positive correlations across the majority of the 22 supermodule classes (Figure 2B). However, no clear pattern of positive and significant correlations was observed between the systems when the underlying results were investigated.

**Fig 2.**
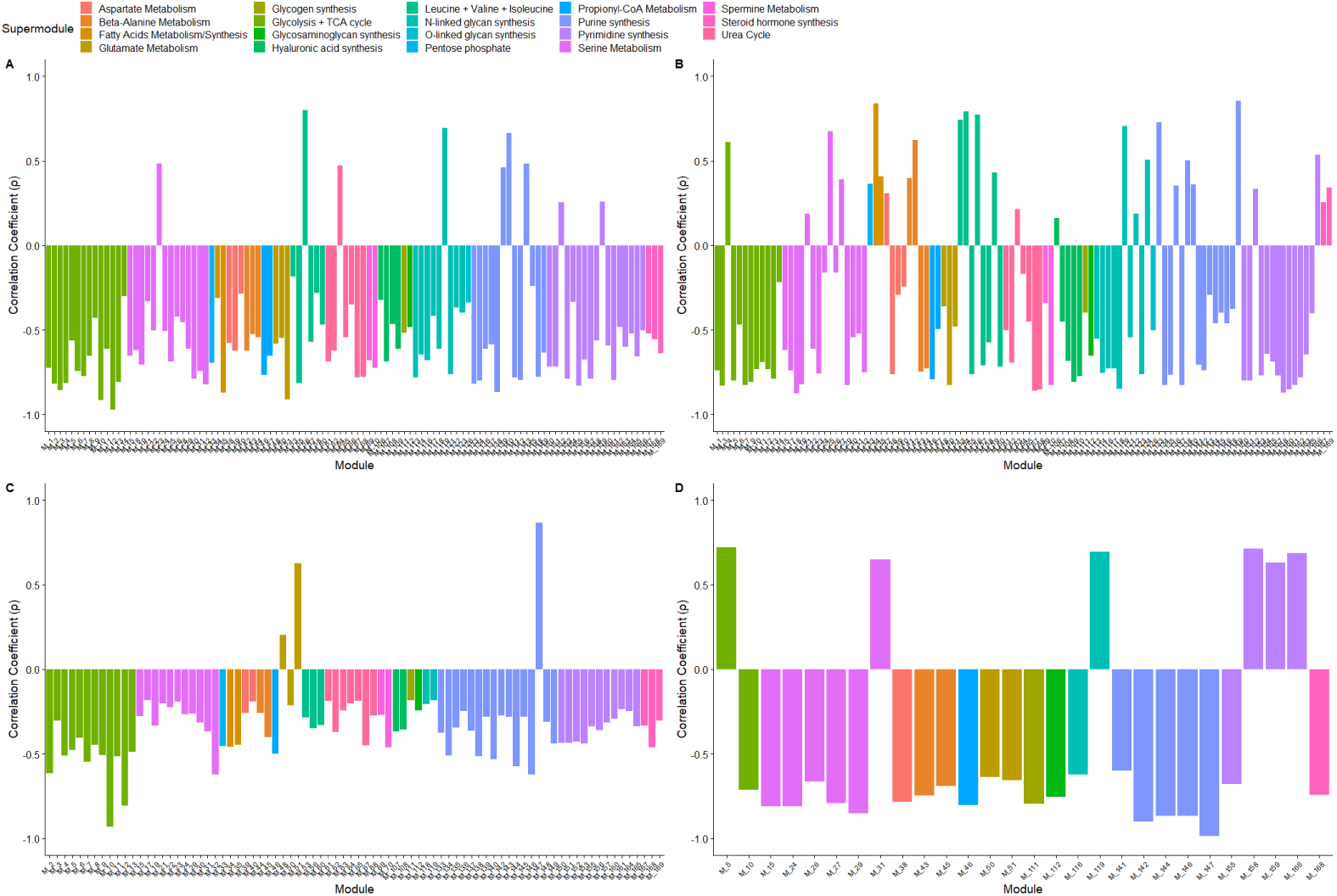
Significant (p *<* 0.05) Spearman correlations between the maximum positive Compass penalty within a collection of reactions approximating those in an scFEA module and the predicted scFEA flux for the analogous module. The results of both systems run on three datasets were used for comparison. (A) A single-cell multiple myeloma dataset. (B) The same multiple myeloma dataset with imputation measures enabled in both systems. (C) A single-cell fetal mouse lung tissue dataset. (D) A bulk ovarian clear cell carcinoma dataset.

Overall, this decomposition of significantly correlating modules into their supermodule classes suggests that scFEA and Compass can achieve consensus within at least a subset of module and supermodule classes using the maximum penalty method. Here, substantial agreement existed within the large supermodule collections of glycolysis and TCA, serine metabolism, and pyrimidine synthesis when run on single-cell datasets without imputation. Consensus among a broad range of supermodule classes was greatest for the multiple myeloma results without imputation, suggesting that a higher consensus may be obtained for a broad range of processes in single-cell data arising from cancers.

### Subsystem Reactions to Module Similarity

The agreement between the systems that was observed at the module level was based on an automated selection of a single reaction penalty. To determine whether the level of agreement observed at the module level also existed at the reaction level, related groups of Recon reaction identifiers corresponding to a biological process (“subsystems”) were matched to their analogous scFEA supermodule classes. This allowed the results from the module-level consensus to be further investigated at the reaction level. Spearman correlations were performed between the Compass forward reaction penalties within the select Recon2 subsystems and the scFEA predicted fluxes for the modules within the analogous supermodule. The subsystems and supermodule classes related to TCA, glycolysis and purine synthesis were selected. Glycolysis and TCA were chosen based on the module-level performance within these groups for the single-cell results, and their importance to cancer cell metabolism. Purine synthesis was selected for further analysis based on the number of unexpectedly positive correlations for the single-cell result comparison at the module level.

Heatmaps of the single-cell results showed significant negative correlations between the majority of the scFEA glycolysis and TCA modules, and the majority of the reaction penalties within the Recon2 TCA or glycolysis subsystems (Figure 3A-3B, 3G-3H). For the multiple myeloma results, the first four scFEA modules, representing the modularized reactions converting glucose to pyruvate, and module six, representing the conversion of pyruate to lactate, showed overall moderate-to-strong negative correlations with all but one glycolysis reaction penalty. Module five (pyruvate to acetyl-coa) showed reduced but significant negative correlations with the same reactions. The mouse lung results showed a weaker, but similar pattern to the multiple myeloma results for modules two to six (Figure 3G). For the multiple myeloma results, significant and negative correlations were observed between the TCA cycle subsystem reactions and the scFEA TCA predicted fluxes, modules seven to fourteen, for all but two reactions (Figure 3B).

**Fig 3.**
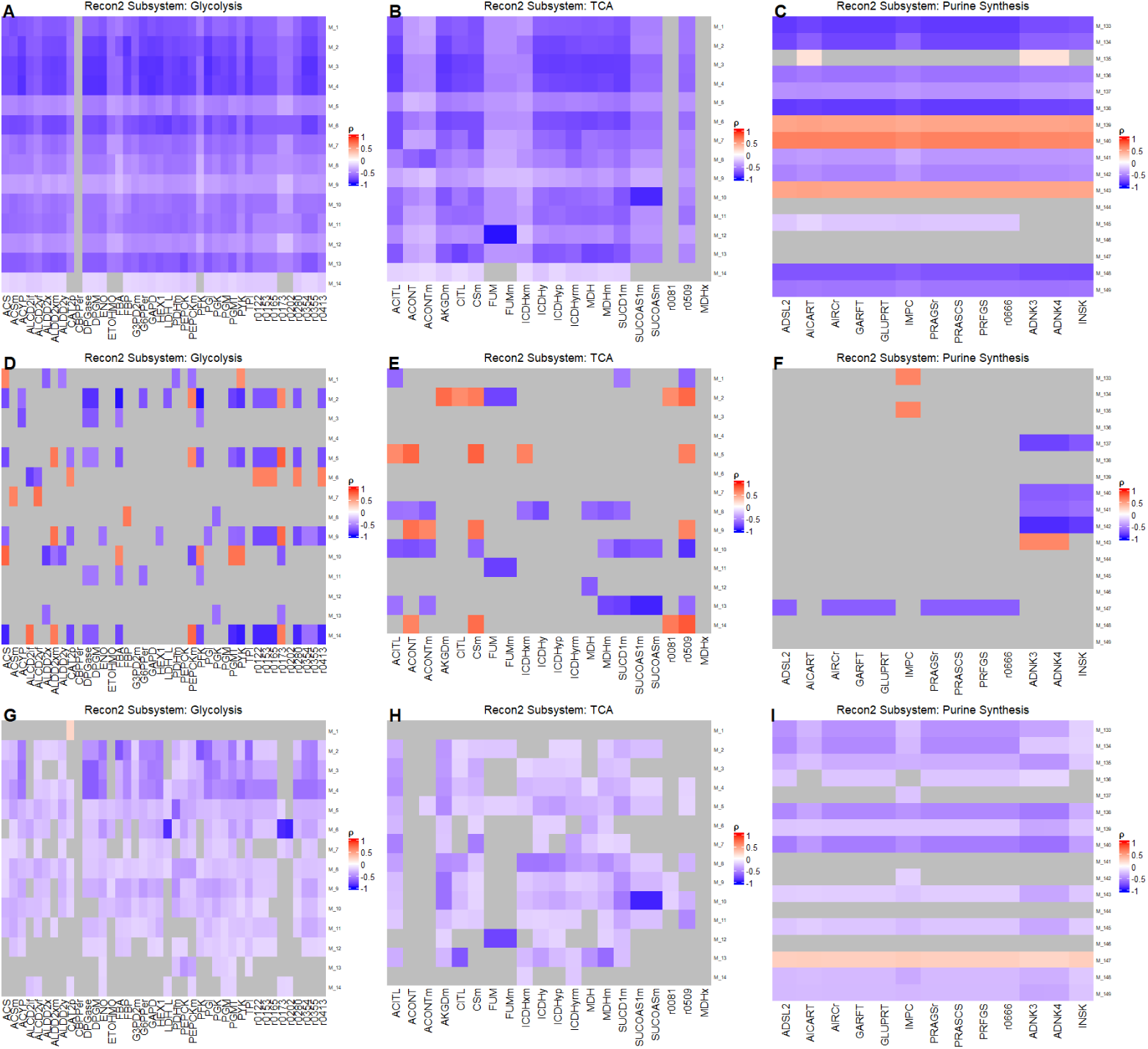
Significant (p *<* 0.05) Spearman correlations between reactions within a specified Recon2 subsystem and the analogous scFEA supermodule for the Compass and scFEA results from three datasets. Three Recon2 subsystems, glycolysis, TCA, and purine synthesis were selected and their reactions correlated with their analogous scFEA modules. (A-C) Single-cell multiple myeloma results (D-F) Bulk ovarian clear cell carcinoma results (G-I) Single-cell mouse fetal lung results.

A reduction in significant correlations was noted generally for the glycolysis and TCA cycle results between the multiple myeloma and mouse results (Figures 3A, 3B, 3G, 3H). No clear patterns in correlation were observed for any of the bulk results (Figures 3D-3F). This reaction-level analysis suggests that the results of scFEA and Compass results can achieve a general consensus at the reaction level for glycolysis and the TCA cycle, particularly in human cancers, by comparing the forward reaction penalties produced by Compass to the scFEA predicted fluxes for analogous collections of modules and reactions. However, the reaction-level consensus performance may be reduced when mouse cell data is used as input, and the results may be non-interpretable when bulk data is used.

To investigate the relationship between the module-level and reaction-level results, the reaction heatmaps were compared to the results based on the maximum module penalties. For the module-level mouse results, two of the fourteen modules, one and fourteen, within the glycolysis and TCA supermodule class, did not demonstrate significant correlations between the maximum positive penalty and the corresponding scFEA flux (Figure 2C). The absence of these modules were reflected by one weak correlation between module one and every glycolysis reaction within the subsystem, or two weak correlations between module fourteen and every reaction in the TCA cycle subsystem (Figure 2C, 3G, 3H). For the module-level multiple myeloma results, each one of the fourteen modules showed significant correlations across every glycolysis and TCA module (Figure 2A). This was reflected in the negative correlations between each module and nearly every reaction in glycolysis or TCA at the reaction-level (Figure 2A, 3A, 3B). Thus, the use of the maximum positive penalty method is supported at the reaction level.

The moderate-to-strong positive correlations for purine synthesis that were observed at the module level were investigated at the reaction-level. For the multiple myeloma and mouse lung results, consistently positive correlations at the reaction level corresponded exactly to the modules with positive correlations in the module-level analysis (Figures 2A, 2C, 3C, 3I). For the mouse results, weak positive correlations were noted between module 147 and each Recon2 purine synthesis reaction penalties. For the multiple myeloma results, moderate-to-strong correlations were observed between the purine synthesis reaction penalties and modules 139, 140 and 143 (Figure 3A, 3C). The underlying scFEA flux predictions for these modules did not demonstrate any consistent pattern to explain this phenomenon. Notably, in addition to the broad pattern of negative correlations across glycolysis and TCA, both of the single-cell results demonstrated significant negative correlations between the forward reaction penalties for “ACITL” and “SUCOAS1m, SUCOASm” and their corresponding scFEA TCA modules, seven and ten respectively, despite the reactions being oppositely directed to their module analogs (Figure 3B, 3H). This suggests that reaction direction does not reduce the significance of a correlation or change its correlation coefficient to a positive value. Thus, consistently positive correlations between reaction penalties and scFEA module fluxes may represent genuine disagreement between the two systems and can be detected by the maximum positive penalty method at the module level.

### scFEA and Compass performance and consensus testing on matched scRNA-seq and metabolomics data

Using the paired metabolomics and transcriptomics data of both Wagner et al. and Alghamdi et al., we tested the predictions produced by both scFEA and Compass against the corresponding metabolomics dataset.

### Consistency of scFEA predicted fluxes with established Th17 metabolomic measurements

To determine the consistency of the scFEA predicted fluxes with the metabolomics data of Wagner et al., scFEA was run on separate pathological and normal Th17 datasets and compared to the metabolomics data for glycolysis, TCA and polyamine metabolism [5]. Of the fourteen TCA and glycolysis modules, thirteen showed highly significant (p*_adj_ <* 0.0001) differences in the means between the normal and pathological cells (Figure 4). The succinate-to-fumarate (module 11) did not produce a significant difference between the cells types. Of the remaining thirteen modules, six could be directly related to the Th17 steady-state and fresh media LC/MS metabolomics results of Wagner et al. The products of modules two, three and six (G3P, 3PD and lactate), were measured by Wagner et al. as one of seven glycolysis metabolites. The products of modules seven, ten and twelve (citrate, succinate and malate) were measured by Wagner et al. as one of six TCA cycle metabolites. Additionally, the glycolytic flux producing G6P (module one) could be indirectly related to the hexose monophosphate measurements of Wagner et al. Consistent with the steady-state metabolomics of Wagner et al., the means of the scFEA predicted fluxes for the TCA cycle, producing citrate, succinate and malate (modules seven, ten and twelve), were significantly elevated by greater than 0.002 units in the pathological versus the normal Th17 cells (Figure 4). Of the glycolysis modules, both modules two and three, producing either G3P or 3PD, showed significant mean increases in the pathological cells. These elevations were consistent with either the “DHAP/G3P” or “3-phosphoglycerate” measurements of Wagner et al. However, only module three demonstrated a substantial mean increase of 0.0037 between the pathological and normal predictions, while module two produced only a marginal mean flux elevation of 4.58 *×* 10*^−^*^6^ between the two cell types. In contrast, both the G6P and lactate producing modules (modules one and six) showed significant and substantial increases of either 0.0021 (module one) or 0.0018 (module six) in the mean predicted fluxes for the normal cells, which were at odds with either the hexose monophosphate or lacate metabolomic measurements of Wagner et al. Although not in the metabolomics data of Wagner et al., four remaining scFEA predictions for the fluxes producing pyruvate, acetyl-coa, 2OG and succinyl-coa were consistent with the general glycolysis and TCA cycle increases reported in the pathological state by the Seahorse experiments of Wagner et al [5]. However, of the two oxaloacetate producing fluxes predicted by scFEA, malate-to-oxaloacetate and pyruvate-to-oxaloacetate, only malate-to-oxaloacetate was predicted to be signficantly increased in the pathological cells. A marginal increase in mean flux was noted for the control Th17 cells for the pyruvate-to-oxaloacetate module.

**Fig 4.**
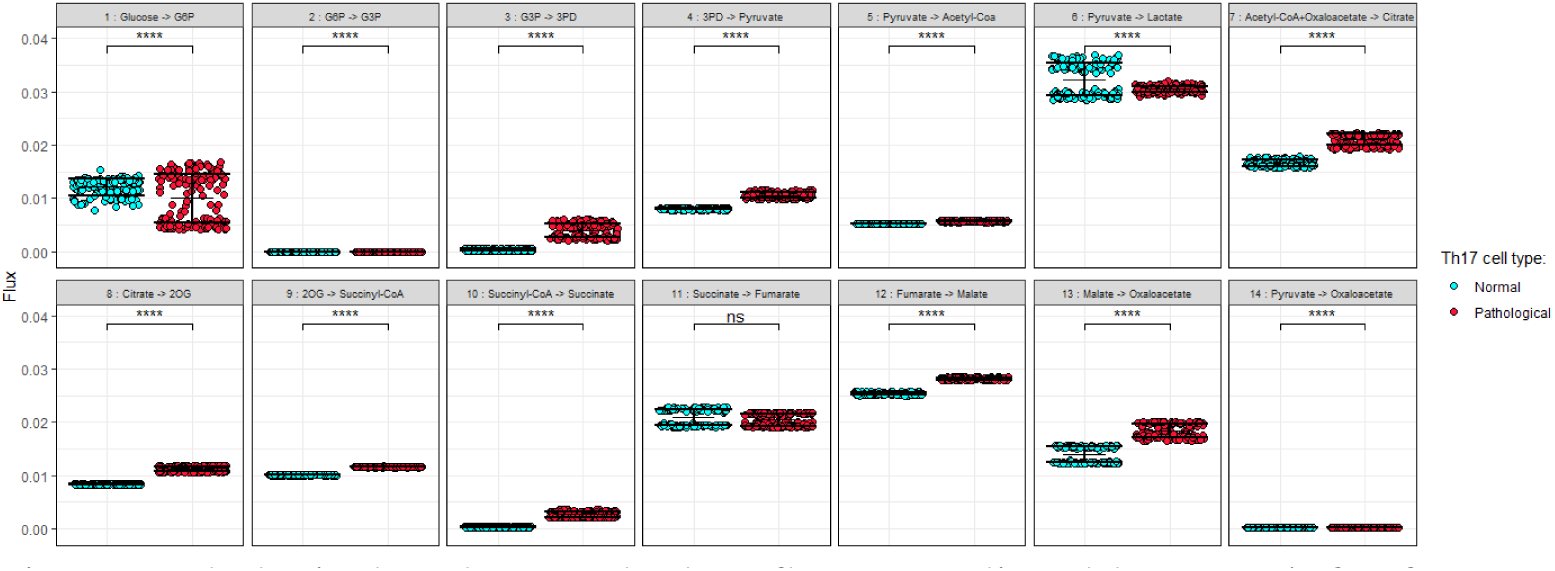
Pathological and normal Th17 fluxes predicted by scFEA for fourteen glycolysis and TCA modules are shown. Pathological cell results are shown in red and non-pathological are shown in blue. The glycolysis modules are modules one-to-six, and the TCA modules are modules seven-to-fourteen. The mean predicted values for each cell class is indicated by the central crossbar and the standard error from the mean is represented by the outer bars. A Welch’s t-test was performed between the predicted fluxes for each module between the pathological and normal cell results.The significance is based on a Bonferroni adjusted p-value. **** Highly significant results (p *<* 0.0001). (ns) non-significant (p *≥* 0.05) results.

The polyamine-related metabolomics data of Wagner et al showed significant increases in creatine for the control cells and n-acteyl-putrescine and putrescine in the pathological cells [5]. Two of the three scFEA predicted fluxes related to these metabolites were consistent with the data of Wagner et al. For the ornithine-to-putrescine module predictions of scFEA, the pathological Th17 cells had a highly significant increase in predicted flux relative to the control cells (Figure 5). The predicted arginine-to-putrescine fluxes were unexpectedly constant across both cell types, although the mean was higher by 3.5 *×* 10*^−^*^4^ units in the pathological cells. However, in contrast to the creatine results in the Compass metabolomics, the predicted mean flux for the glycine-to-creatine module was 3.5 *×* 10*^−^*^3^ units higher in the pathological cells.

**Fig 5.**
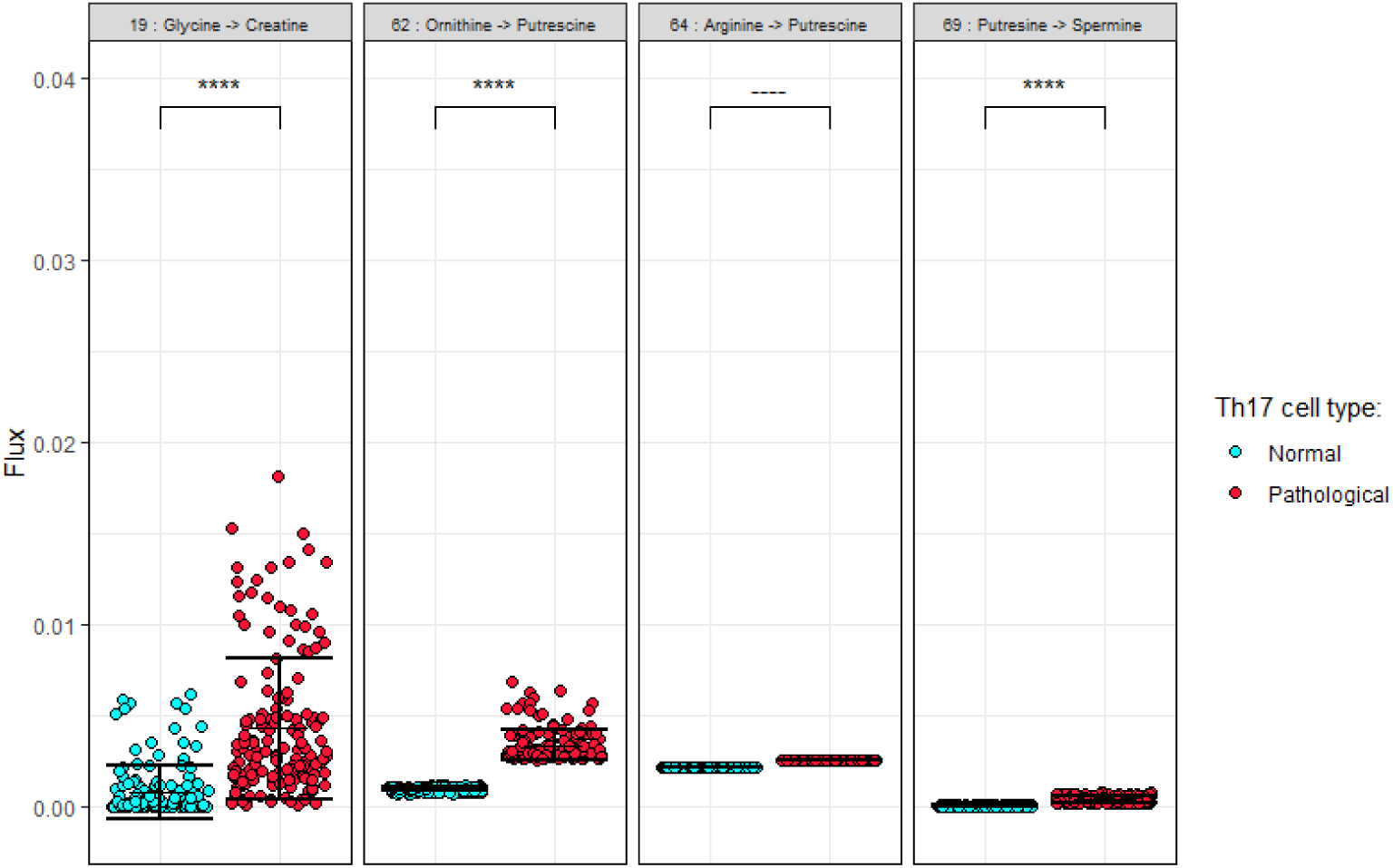
Four fluxes scFEA predicted fluxes for polyamine-related metabolism for pathological and normal Th17 cells. The fluxes for the pathological cells are indicated by red, and the normal cells are indicated by blue. The mean values are indicated by the crossbar and the standard deviation from the mean is given by the upper and lower bars in each plot. **** Highly significant results (p *<* 0.0001). ---- Constant values for the predicted pathological and normal fluxes for both cell types for which no adjusted p-value could be calculated.

Collectively, the scFEA results for the TCA and polyamine modules provide partial support for the hypothesis that scFEA can predict mouse Th17 cell metabolism that is consistent with experimentally-measured metabolic properties. However, this was contradicted in the case of glycolysis.

### Compass Th17 results

The post-processing code of Wagner et al. was adapted to approximate their analysis with the results from two of our runs using segregated pathological and normal datasets. Similar results to Wagner et al. were obtained for glycolysis, and the predicted Cohen’s distance was similar across the labeled reactions, with general and significant increases in the majority of glycolytic reactions in the pathological cells (Figure S1, top left). For the labeled TCA cycle reactions, aconitate hydratase, the reaction with the lowest adjusted p-value in the analogous figure of Wagner et al., was not significantly elevated in the pathological cells (Figure S1, top right). Notably, in contrast to the results of Wagner et al., the reaction penalties for putrescine diamine oxidase, within amino acid metabolism, were identical and constant across both the pathological and normal cells.

These results suggest that segregating the pathological and normal Th17 data when k-nearest neighbors smoothing is enabled, or the use of FPKM versus TPM counts, can affect the Compass results for select outlier reactions. With these data, Compass achieved the results most consistent with the metabolomics and Seahorse assays of

Wagner et al. for glycolysis [5]. This was in contrast to scFEA, which showed a higher consistency with the metabolomics for TCA. Collectively, this suggests that scFEA and Compass disagree on the predicted differences for glycolysis and TCA between pathological and normal Th17 cells, but each system is partially consistent with the direct metabolic measurements.

### Consensus measures between scFEA and Compass Th17 predictions

To determine whether the fold-change correlations for each of the two systems reflected a consensus between the systems at the cell or module level, the maximum positive penalty consensus tests from sections “Automating the Comparison of Compass and scFEA results” and “Comparison of modularized Compass penalties to scFEA predicted fluxes using three datasets” were applied to the APEX1-KD and control Pa03c results from both scFEA and Compass. For the Compass predictions, k-nearest neighbors smoothing was enabled and disabled to determine its effect on the consistency between the systems. For both the pathological and normal predictions, weak but highly significant negative cell-wise correlations were observed between the maximum positive compass penalties per-cell and the predicted scFEA fluxes (Table 2).

**Table 2.** The cell-wise and module-wise consensus measures for the flux predictions of both scFEA and Compass when run on scRNA-seq data from normal (“Th17n”) and pathological (“Th17p”) *Mus musculus* Th17 cells. The predictions of Compass with both KNN smoothing enabled and disabled were correlated with the corresponding scFEA flux predictions. The maximum positive module penalty was used. Spearman correlation coefficients were averaged after Fisher’s Z was applied and back-transformed following the calculation. The number of modules with a significant (p *<* 0.05) and positive correlation is indicated by positive *ρ*. The ratio of significant and positively correlated modules is indicated in the sixth column.

| Th17n with KNN smoothing |  |  |  |  |  |
| --- | --- | --- | --- | --- | --- |
| Cell |  | Module |  |  |  |
| r | $p_{comb}$ | $p < 0.05$ | NA | Positive $\rho$ | $\frac{\text{Positive } \rho}{p < 0.05}$ |
| -0.30 | $9.5 \times 10^{-302}$ | 48 | 19 | 26 | 0.54 |
| Th17n without KNN smoothing |  |  |  |  |  |
| Cell |  | Module |  |  |  |
| r | $p_{comb}$ | $p < 0.05$ | NA | Positive $\rho$ | $\frac{\text{Positive } \rho}{p < 0.05}$ |
| -0.30 | $4.9 \times 10^{-300}$ | 44 | 19 | 25 | 0.57 |
| Th17p with KNN smoothing |  |  |  |  |  |
| Cell |  | Module |  |  |  |
| r | $p_{comb}$ | $p < 0.05$ | NA | Positive $\rho$ | $\frac{\text{Positive } \rho}{p < 0.05}$ |
| -0.30 | $1.1 \times 10^{-272}$ | 71 | 16 | 30 | 0.47 |
| Th17p without KNN smoothing |  |  |  |  |  |
| Cell |  | Module |  |  |  |
| r | $p_{comb}$ | $p < 0.05$ | NA | Positive $\rho$ | $\frac{\text{Positive } \rho}{p < 0.05}$ |
| -0.30 | $1.3 \times 10^{-270}$ | 65 | 16 | 30 | 0.46 |

A clear increase was observed in the number of significantly correlated modules for the pathological Th17 cells, as well as a more favorable ratio of positive-to-significant correlations (Table 2). Enabling KNN smoothing in Compass slightly increased the number of significantly correlated modules for both cell types, but had a negligible effect on the ratio of positive to total significantly correlated modules. These results support the idea that metabolic dysregulation in pathological Th17 cells increases the general module-wise consensus, but not the cell-wise consensus, relative to the control.

The modules for glycolysis, TCA, and polyamine metabolism were looked at in isolation to determine the level of agreement between the results for Compass and scFEA. For the normal Th17 cells, three of the fourteen glycolysis and TCA modules, and no polyamine modules, showed a significant negative correlation between the Compass maximum positive module penalty and the scFEA-predicted flux (Figures 6A-D). For the pathological Th17 cells, half of the fourteen glycolysis and TCA modules, and one polyamine module, showed significant negative correlations when KNN smoothing was disabled (Figure 6D). Of the glycolysis and TCA modules, one, five and ten were significantly correlated across both the pathological and normal Th17 datasets, with KNN smoothing enabled and disabled (Figures 6A, 6B). Notably, the strongest correlation between the two systems was found across all datasets for module ten, succinyl-coa-to-succinate, which was the most significantly increased TCA cycle reaction predicted by Compass (Figures 6A-6D, S1). No clear pattern for the polyamine modules was observed across the dataset and setting combinations. These results imply that the disagreement between the systems that was observed for glycolysis and TCA in the previous Compass and scFEA results (sections “Compass Th17 results” and “Consistency of scFEA predicted fluxes with established Th17 metabolomic measurements”), and between the systems and the metabolomics, may be reflected in the module-wise consensus.

**Fig 6.**
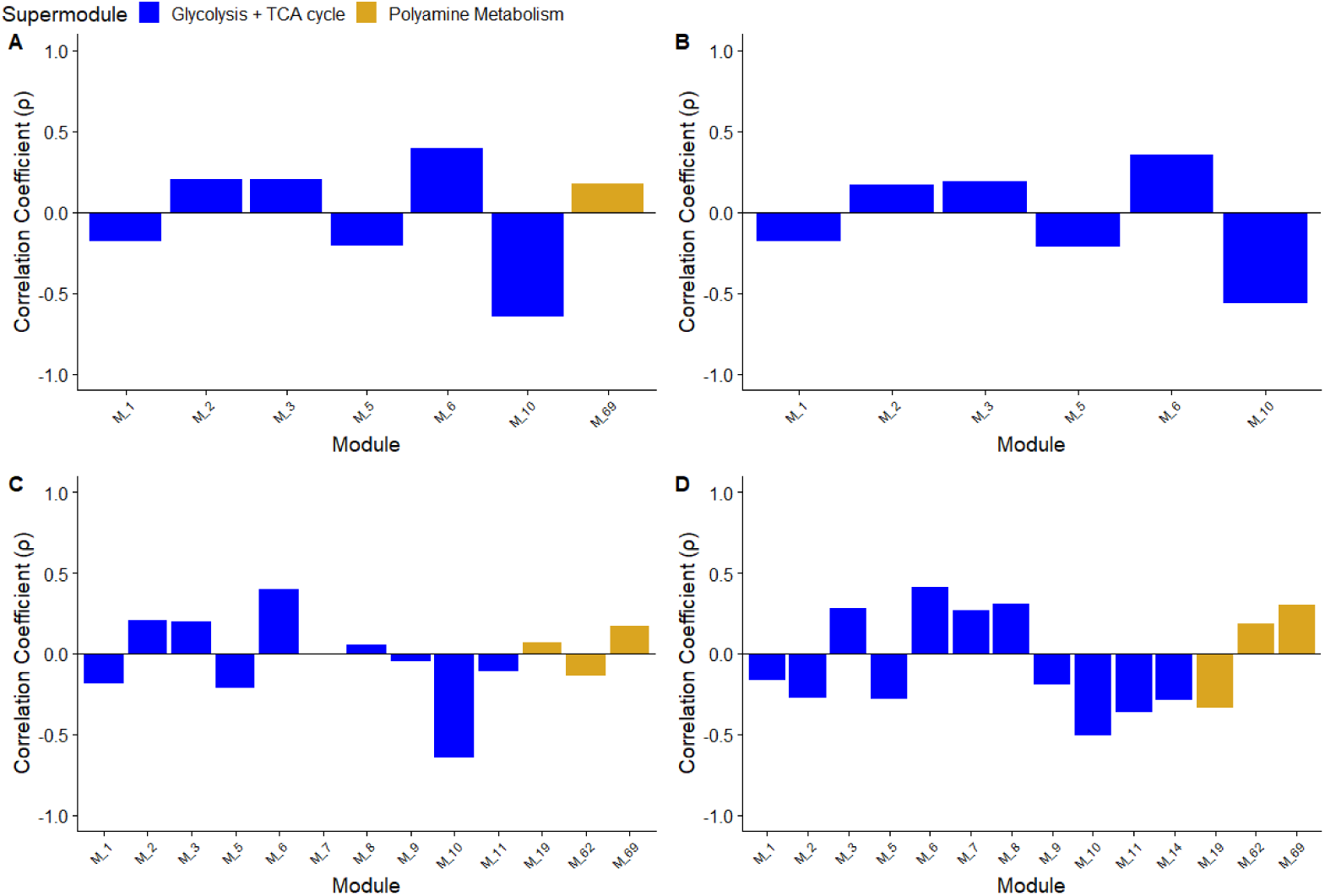
Module-wise consensus measures applied to the Compass and scFEA predictions for pathological and normal Th17 cells. The maximum Compass penalty associated with a forward reaction in a cluster of reactions approximating those in an scFEA module were correlated with the scFEA predicted flux for same module, for 131 non-transport modules. Modules that showed a significant (p *<* 0.05) Spearman correlation between a Compass derived module penalty measure and the corresponding scFEA module were selected for inclusion. The plots are colored according to either the glycolysis and TCA supermodule class used by scFEA, or by membership in one of the four polyamine-related modules. (A-B) normal Th17 cells and (C-D) pathological Th17 cells. (B-D) results with Compass KNN smoothing disabled.

### Compass predictions on scFEA data

Alghamdi et al. validated scFEA by running the system on two sets of pancreatic adenocarcinoma (“Pa03c”) cells for which they had generated matched gene expression and bulk metabolite abundances [1]. The authors validated their glycolysis and TCA predictions by performing a Pearson correlation between the mean predicted flux fold changes and their corresponding measured metabolic product fold changes [1].

To measure the similarity between the predicted activities and the metabolite abundances of Alghamdi et al., the directed reactions in Compass’ metadata were manually mapped to the glycolysis and TCA cycle intermediates in the scFEA meta-bolomics data. Eleven reactions, including seven cytoplasmic variants and four mitochondrial variants, were mapped to a product compound (Table 3). A subset of six reaction and metabolite pairs was found to be comparable to the predicted flux versus metabolite fold change results of Alghamdi et al. (Table 3). A seventh metabolite, 2-oxoglutarate (2OG), was mapped to a reaction and added to the Compass metabolite to reaction map, as oxaloacetic acid (OAA) could not be located in the scFEA metabolomics file.

**Table 3.**
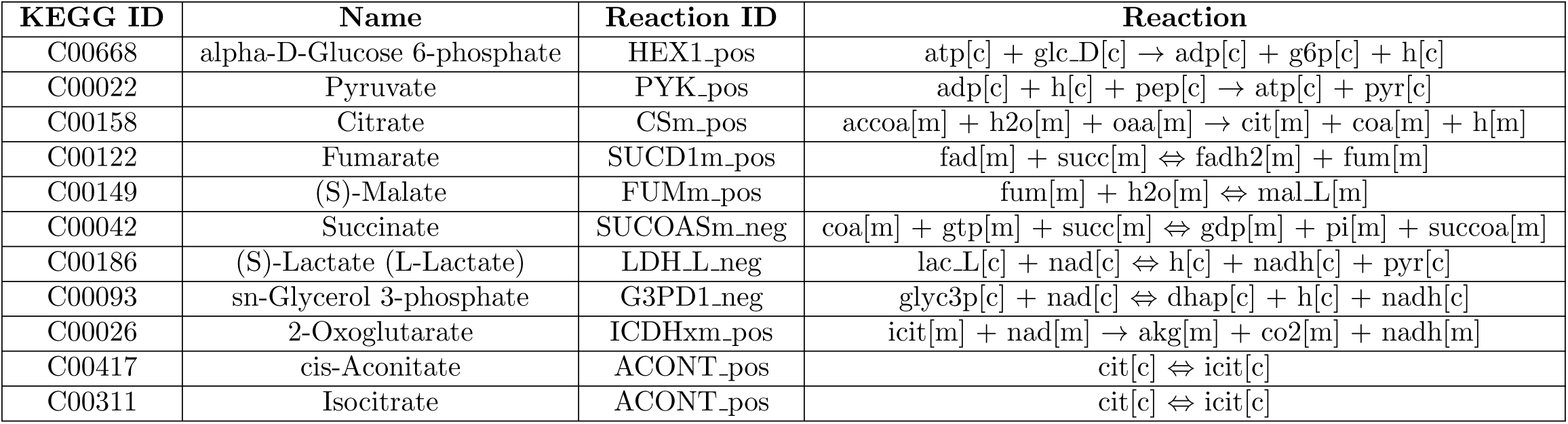
Glycolysis and TCA compounds mapped to Recon reactions used to compare Compass Pa03c results to the metabolomics data from Alghamdi et al. The KEGG compound identifier, compound name and directed Recon identifier, used by Compass, are listed in columns 1-3. The reaction identifier and its associated reaction were obtained from the Virtual Metabolic Human (’VMH’) database. Cytoplasmic reactions are distinguished from mitochondrial variants by “c” or “m” associated with each reactant or product, and by the lowercase “m” in the Recon identifier. The first six compounds (alpha-D-Glucose 6-phosphate to succinate) were used in a similar analysis by Alghamdi et al. The final five compounds were in the metabolomics data of Alghamdi et al. and matched to a Compass reaction.

| KEGG ID | Name | Reaction ID | Reaction |
| --- | --- | --- | --- |
| C00668 | alpha-D-Glucose 6-phosphate | HEX1_pos | atp[c] + glc_D[c] → adp[c] + g6p[c] + h[c] |
| C00022 | Pyruvate | PYK_pos | adp[c] + h[c] + pep[c] → atp[c] + pyr[c] |
| C00158 | Citrate | CSm_pos | accoa[m] + h2o[m] + oaa[m] → cit[m] + coa[m] + h[m] |
| C00122 | Fumarate | SUCD1m_pos | fad[m] + succ[m] ⇌ fadh2[m] + fum[m] |
| C00149 | (S)-Malate | FUMm_pos | fum[m] + h2o[m] ⇌ mal_L[m] |
| C00042 | Succinate | SUCOASm_neg | coa[m] + gtp[m] + succ[m] ⇌ gdp[m] + pi[m] + succoa[m] |
| C00186 | (S)-Lactate (L-Lactate) | LDH_L_neg | lac_L[c] + nad[c] ⇌ h[c] + nadh[c] + pyr[c] |
| C00093 | sn-Glycerol 3-phosphate | G3PD1_neg | glyc3p[c] + nad[c] ⇌ dhap[c] + h[c] + nadh[c] |
| C00026 | 2-Oxoglutarate | ICDHxm_pos | icit[m] + nad[m] → akg[m] + co2[m] + nadh[m] |
| C00417 | cis-Aconitate | ACONT_pos | cit[c] ⇌ icit[c] |
| C00311 | Isocitrate | ACONT_pos | cit[c] ⇌ icit[c] |

Compass was run on either the control or APEX1-KD Pa03c data with KNN smoothing enabled or disabled. For each result, the reactions mapping to each of the eleven metabolites were extracted, and an activity measure was calculated for the reaction by taking the inverse of the reaction penalty.

Between the 48 control and 48 APEX1-KD Pa03c cells, the mean fold change in each reaction activity and the mean fold change in its corresponding metabolite were correlated by the Pearson method. For the productive reactions, the Pearson correlation coefficient was 0.31 for the seven-metabolite subset (p=0.50) without imputation and 0.33 (p=0.47) with imputation (Figure 7). The eleven metabolites and their associated reactions showed a correlation coefficient of 0.36 (p=0.28) without KNN smoothing and 0.37 (p=0.26) with KNN enabled. KNN smoothing produced only a marginal difference in the results for these samples. Overall, the results suggest that a sufficiently large number of predicted Compass reaction activities could provide a limited coherence with direct metabolomics measurements when considering the glycolysis and TCA cycle flux alterations that occur as a result of an APEX1 knockdown in Pa03c cells.

**Fig 7.**
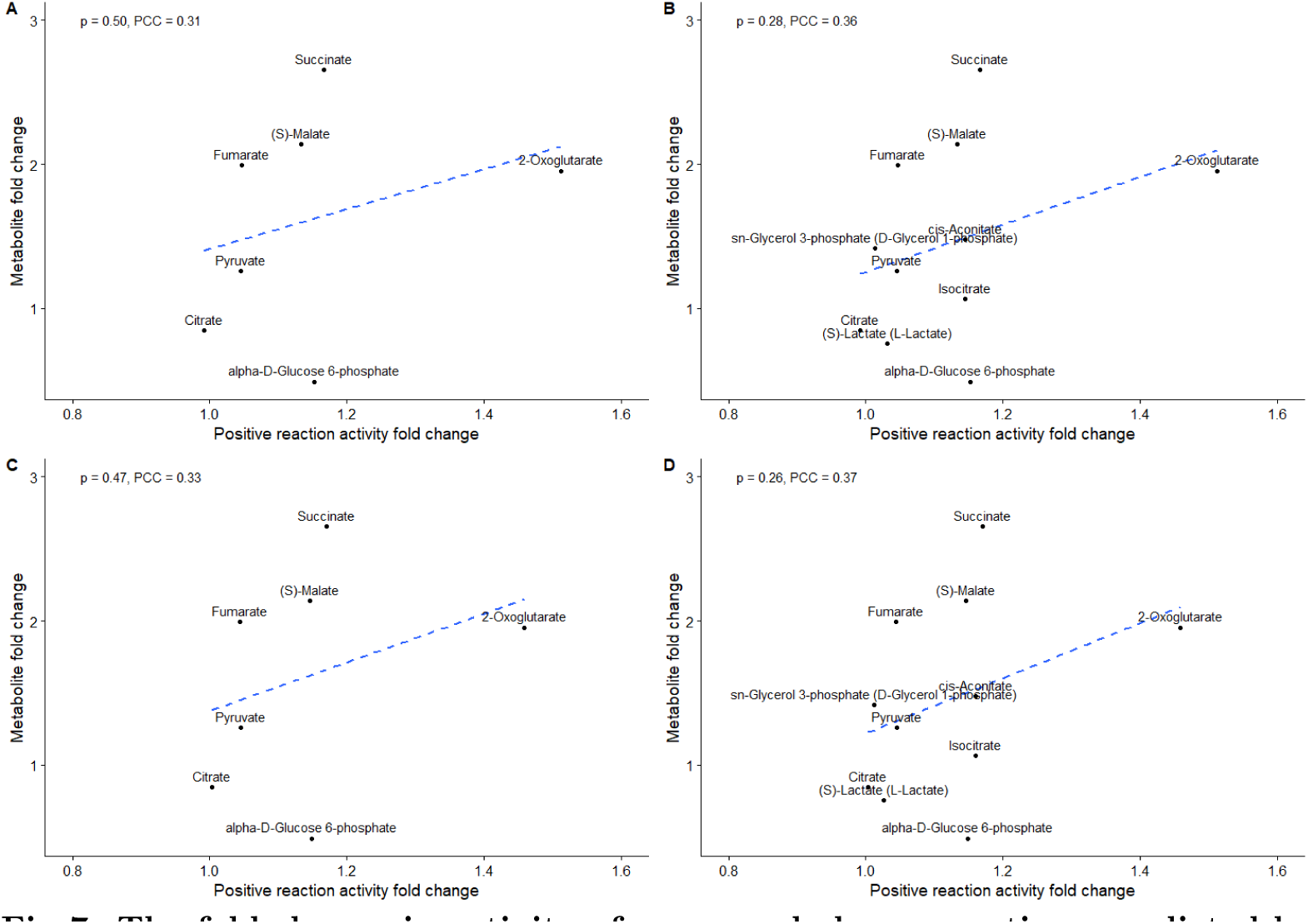
The fold change in activity of seven and eleven reactions predicted by Compass between control and APEX1 knockdown Pa03c cells versus the corresponding change in metabolite produced by each reaction. (A,C) Seven reactions and their metabolic products. (B,D) Eleven reactions and their metabolic products. (A,B) Results without KNN smoothing enabled. (C,D) Results with KNN smoothing enabled. A linear model calculated from the results is shown in blue. The Pearson correlation coefficient (“PCC”) and p-value are indicated in the top left of each figure.

### scFEA Pa03c testing

As a reference measure, we ran scFEA on the same 48 APEX1-KD and 48 control Pa03c cell datasets as Compass. Similar to the Compass analysis, each scFEA module was mapped to a TCA or glycolysis metabolite. The fold changes in mean flux and the fold changes in the corresponding metabolites were calculated between the control cells and the APEX1-KD cells. The flux and metabolite fold changes were correlated by the Pearson method. As a module producing glucose-1-phosphate (G1P) could not be determined, we omitted it from our analysis. For OAA, we substituted in 2OG and its corresponding module, similar to the Compass analysis.

The initial results showed a Pearson correlation coefficient of -0.18 (p=0.69) (Figure 8A). A large outlier was observed for the 3PD to pyruvate module. The predicted flux through this module was nearly 40 fold higher in the control versus the APEX1-KD cells. Removal of the outlier resulted in a Pearson correlation coefficient of 0.54 (p=0.27), but did not produce a significant correlation coefficient (Figure 8B).

**Fig 8.**
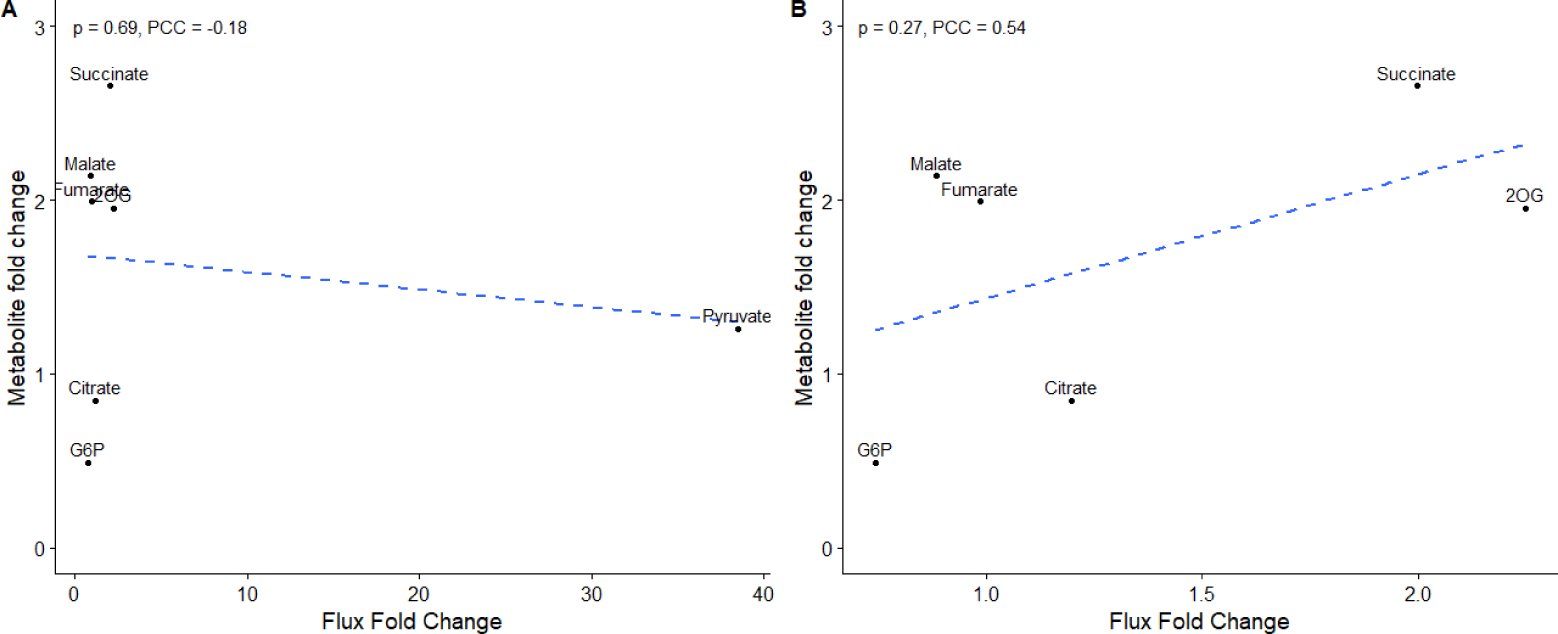
The scFEA predicted fold change between Pa03c control and APEX1 knockdown cells in the fluxes through seven modules versus the corresponding change in their experimentally measured product metabolites. A linear model is shown in blue and the Pearson Correlation Coefficient (“PCC”) and p-value are shown in the top left of each panel. (A) With all seven metabolites. (B) With the pyruvate outlier removed.

Our results vary from those of the 27 APEX1-KD and 46 control cells of Alghamdi et al. However, our results suggest that scFEA may provide a better ability to predict flux alterations in glycolysis and TCA cycle metabolism when using unfiltered pancreatic adenocarcinoma cell data and when clear outlier results are removed.

### Consensus testing between the Compass and scFEA predictions for Pa03c control and APEX1 knockdown cells

To test the agreement between scFEA and Compass for the two types of Pa03c cells, we used both the cell-wise and module-wise consensus tests and the maximum module positive penalty method from sections “Automating the Comparison of Compass and scFEA results” and “Comparison of modularized Compass penalties to scFEA predicted fluxes using three datasets”. The consensus measures were applied to the APEX1-KD and control Pa03c results from running scFEA and Compass with Compass KNN smoothing enabled and disabled. The significantly correlated glycolysis and TCA modules were plotted for each Pa03c cell-type with and without KNN (Figure 9).

**Fig 9.**
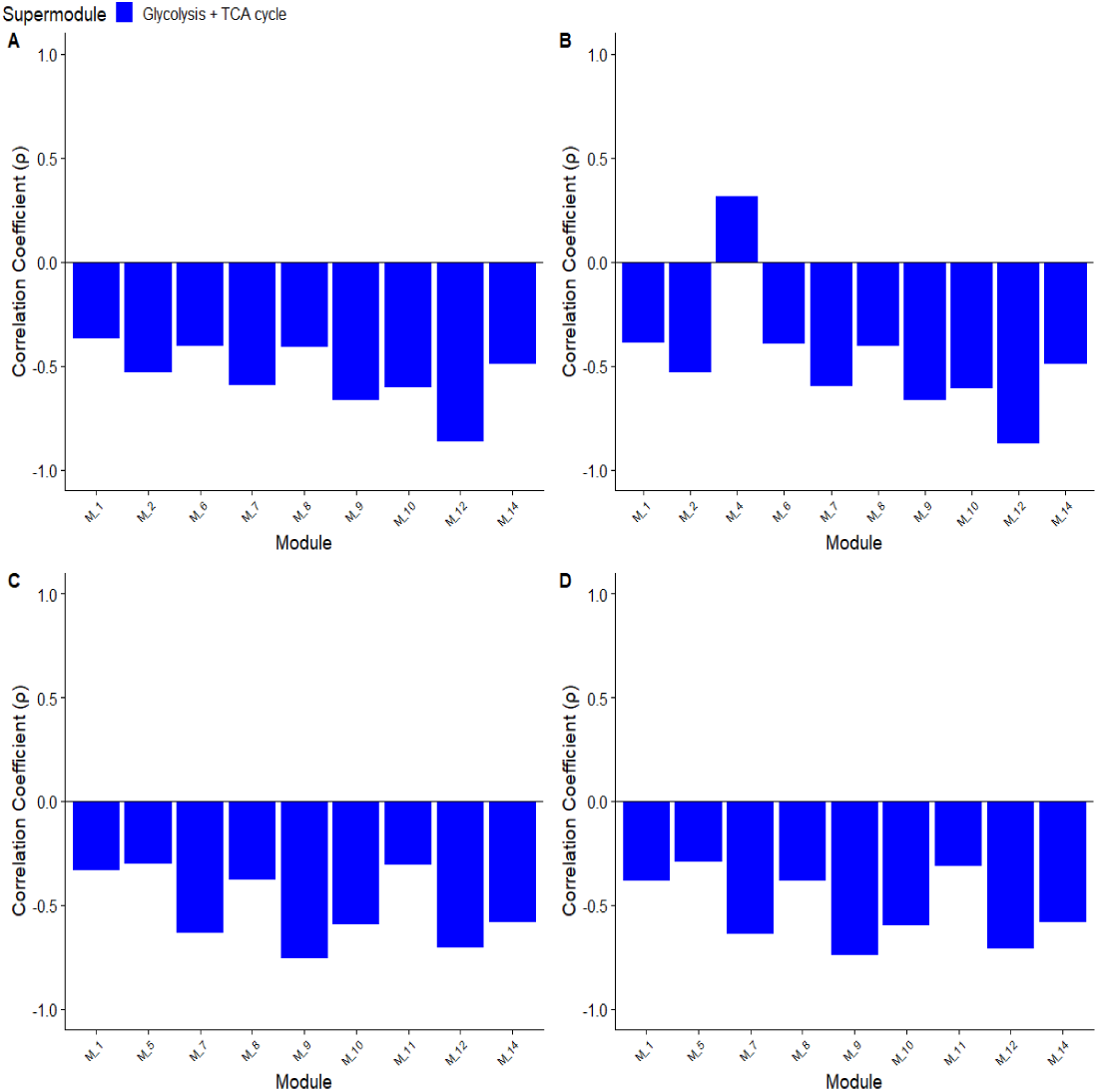
The maximum positive penalty consensus measure applied module-wise to the APEX1-KD and control Pa03c cells. (A-B) The results for the APEX1-KD cells with (A) KNN smoothing enabled in Compass (B) KNN smoothing disabled. (C-D) The results for the control cells (C) KNN smoothing enabled and (D) KNN smoothing disabled.

The cell-wise results showed consistently significant and negative correlation coefficient for each of the cell types and KNN settings. Of these, the APEX1-KD cell-wise correlations showed a combined p-value that was greater than 10^18^ times higher than the control Pa03c cells (Table 4). For the module-wise results, both combinations of cell types and KNN options demonstrated significant correlations for over 63 percent of the 131 total modules. The control cells showed an increase in significantly correlated modules, from 83 in the APEX1-KD cells to 87 in the control. However, three additional unexpected correlations were present for the control. Broadly, this suggests that Compass and scFEA can achieve a modest cell-wise and moderate module-wise consensus when run on single-cell RNA-seq data from pancreatic adenocarcinoma cells. Further, this result suggests that the APEX1 knockdown decreases the cell-wise consensus and module-wise consensus between the systems.

**Table 4.**
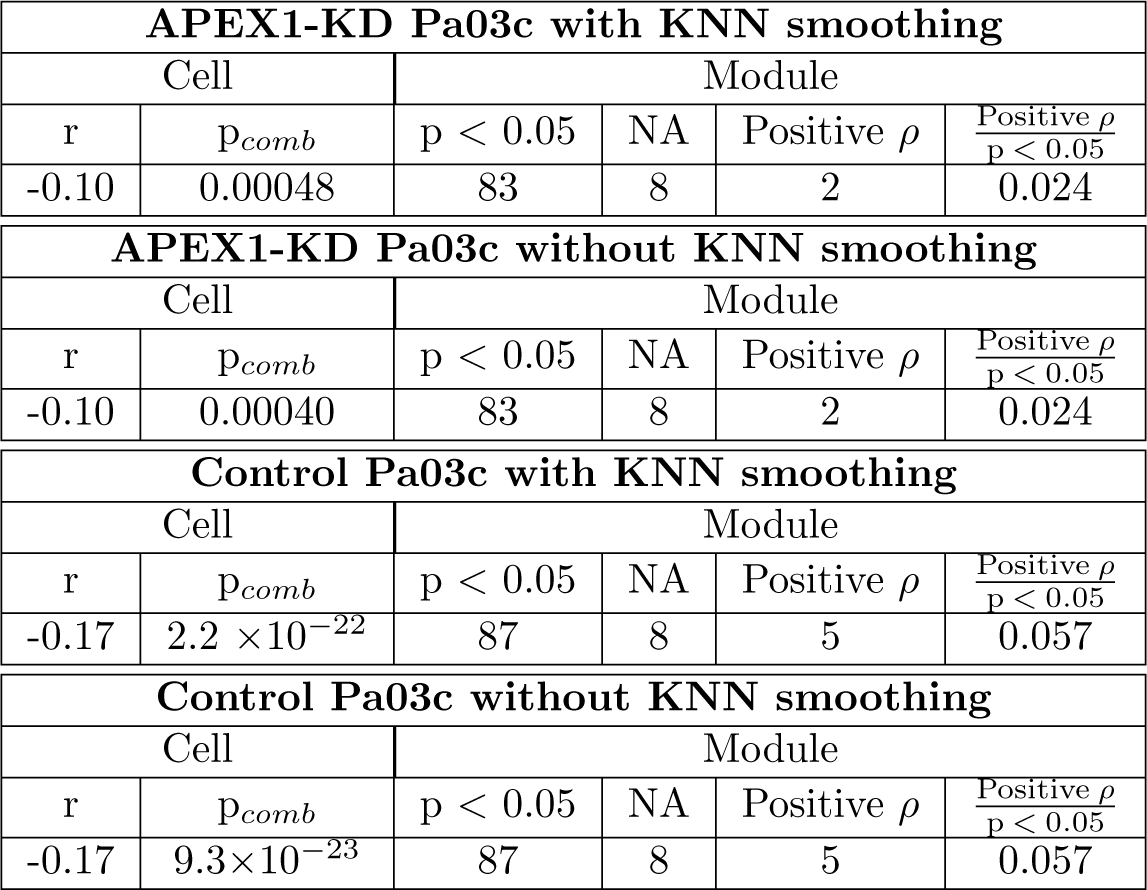
Module-wise and cell-wise consensus measures applied to the Compass and scFEA results for single-cell sequencing data arising from two sets of human pancreatic adenocarinoma cells. The fluxes for the control cells and cells with an apurine endonuclease knockdown (“APEX1-KD”) were predicted by Compass, grouped into scFEA module analogs, and the maximum positive penalty was selected for comparison with the predicted scFEA fluxes. The Spearman correlation coefficient was calculated between the scFEA and Compass results for each cell, and the p-values were combined using Fisher’s method. Results where a static flux prediction across all modules made a correlation impossible are listed as NA. Unexpectedly positive and significant correlations were tallied, and the ratio of such correlations to the total significantly correlated modules are shown in the righthand column.

| APEX1-KD Pa03c with KNN smoothing |  |  |  |  |  |
| --- | --- | --- | --- | --- | --- |
| Cell |  | Module |  |  |  |
| r | $p_{comb}$ | $p < 0.05$ | NA | Positive $\rho$ | $\frac{\text{Positive } \rho}{p < 0.05}$ |
| -0.10 | 0.00048 | 83 | 8 | 2 | 0.024 |
| APEX1-KD Pa03c without KNN smoothing |  |  |  |  |  |
| Cell |  | Module |  |  |  |
| r | $p_{comb}$ | $p < 0.05$ | NA | Positive $\rho$ | $\frac{\text{Positive } \rho}{p < 0.05}$ |
| -0.10 | 0.00040 | 83 | 8 | 2 | 0.024 |
| Control Pa03c with KNN smoothing |  |  |  |  |  |
| Cell |  | Module |  |  |  |
| r | $p_{comb}$ | $p < 0.05$ | NA | Positive $\rho$ | $\frac{\text{Positive } \rho}{p < 0.05}$ |
| -0.17 | $2.2 \times 10^{-22}$ | 87 | 8 | 5 | 0.057 |
| Control Pa03c without KNN smoothing |  |  |  |  |  |
| Cell |  | Module |  |  |  |
| r | $p_{comb}$ | $p < 0.05$ | NA | Positive $\rho$ | $\frac{\text{Positive } \rho}{p < 0.05}$ |
| -0.17 | $9.3 \times 10^{-23}$ | 87 | 8 | 5 | 0.057 |

KNN smoothing had a negligible effect for these consensus measures when implemented in Compass. No effect was noticed when smoothing was enabled for the module-wise measures (Table 4). For the cell-wise control measures, enabling KNN in Compass increased the combined p-value by an order of magnitude (Table 4). Thus, only a minimal effect was observed from Compass imputation when applied to these two datasets.

When plotted, the TCA modules showed a broad agreement between the Compass and scFEA results. At least six of the eight TCA modules, modules seven to fourteen, showed significant negative correlations across each combination of APEX1-KD or control cell with KNN smoothing enabled or disabled in Compass (Figure 9). In each case, at least half of the TCA modules showed a correlation coefficient of less than -0.50. For the glycolysis modules (one through six) three modules showed a significant negative correlation between the Compass module penalty and the predicted scFEA flux in the APEX1-KD results. This was reduced to two in the control results. Of the glycolysis modules, only module two showed a negative correlation of less than -0.50 for the APEX1-KD cells (Figure 9). Notably, module four, 3PD to Pyruvate, which was associated with the 40 fold increase between the control and the APEX1-KD cells, was found to either not correlate, or showed a positive correlation between the Compass module penalty and the scFEA flux. Collectively, these results demonstrate that moderate agreement between the systems can be achieved for TCA, but not glycolysis in Pa03c cells. This agreement is consistent with the independent weak-to-moderate correlations between the systems and the metabolomics measurements that were observed in sections “Compass predictions on scFEA data” and “scFEA Pa03c testing”. Additionally, the presence of non-correlating or positively correlating results for module four supports the use of positive correlations as a measure of disagreement.

## Discussion

Direct measurements of the single-cell metabolome are currently infeasible. Accurate predictions of single-cell flux could provide a high resolution understanding of the altered metabolic states that contribute to pathology, such as in tumor cells, tumor-associated immune cells, and autoimmune cells. These predictions could potentially aid in the discovery of drug targets. However, the methods and outputs of current single-cell flux prediction tools differ greatly. Nonetheless, both scFEA and Compass have been validated by directly measured metabolite data. This suggests that they could be used in concert to provide higher confidence flux estimates, and, if in agreement, to investigate metabolic perturbation at the level of reaction potential. Despite this, comparisons between the performance or output similarities of any transcriptome-based single-cell flux prediction tools do not appear to have been reported.

Here, we showed that Compass results can be grouped in a manner approximating an scFEA module, and that this can be used to compare the similarities in the results of the two systems. Bulk and imputed data caused an increased consensus between the systems for a particular cell, but a reduced number of single modules achieving consensus when every cell was considered. For single-cell data, the module-level agreement that was observed between the two systems when a representative reaction was selected is largely mirrored among related reactions. Our findings suggest that intra-module disagreement between the systems can be divided into two classes: uncorrelated reactions and positive correlations associated with a subsystem-wide disagreement. We found an increased level of consensus between Compass and scFEA for human tumor and mouse autoimmune cells versus healthy mouse cells. Significant agreement across a broad range of processes, including cellular energy production, amino acid metabolism and nucleotide synthesis, was observed. Both scFEA and Compass demonstrated partial consistency with the metabolomics used to validate the opposing system. The disagreement between the systems appeared to reflected in our consensus measures.

### Creation of a Compass-to-scFEA key

Compass and scFEA make use of different metabolic models and reaction identifiers. To determine the feasibility of mapping scFEA reactions to Compass reactions, we were able to generate a key of Compass reaction clusters approximating scFEA modules. At least half the number of reactions as in the original module were present in three-quarters of the modularized groupings. In our testing, the reactions appeared to be correctly mapped to their respective modules, although our reaction clusters contained compartmental or substrate variants. In our limited reaction-level testing, however, the Compass results for compartmental variants appeared to show only minimal differences from their compartment-appropriate counterparts when they were associated with a penalty estimate for a specific dataset. While scFEA and Compass use different reaction identifiers and metabolic models, our results suggest enough overlap between the identifiers to allow for a meaningful comparison between the two systems. This may be due to key metabolic reactions being largely captured by our mapping methods.

### Optimal Compass penalty composites to represent single modules

We sought to determine the degree to which the reactions of Compass could be correlated with an scFEA module. To do so, we needed to select a single Compass penalty, or composite of penalties, to represent a module. Because Compass outputs are at the reaction level, and can be either bidirectional or unidirectional, the process of determining a representative Compass measure for each set of modularized reactions was challenging. Further, the selection of a single penalty measure needed to be agnostic to pathway structure.

To address this, we compared several per-module penalty measures. We found that the best method to compare each of these modularized collections of Compass results to a single scFEA flux was to take the maximum positive penalty per module. We had expected that the forward-to-reverse penalty ratio would provide a more accurate comparison measure, as it could capture net movement through a reaction [20]. However, the maximum positive penalty method showed the strongest single-cell module-wise performance, as measured by the ratio of significant-to-positive correlations. Our results suggest that the forward and reverse penalties are not related to each other in the same manner as kinetic rate constants, or forward and reverse flux ratios, which was our initial assumption [20]. Nonetheless, both the maximum penalty ratio and positive penalties can be used to compare Compass results to scFEA fluxes across modules, with the maximum positive penalty showing the most robust module-wise performance.

### Cell-wise and module-wise consensus for bulk and single-cell data

Although our comparison methods demonstrated strong correlations between the scFEA and Compass results across the majority of modules for our single-cell data, our hypothesis that the systems could be compared was partially contradicted by the cell-wise results. We found that the correlations between the selected penalties for each reaction, and their analogous scFEA module fluxes, were generally poor across individual cells for single-cell data. Nonetheless, in the single-cell results, the correlation coefficients were generally negative, as expected, and the maximum positive penalty method showed significant or nearly significant results. The difference between the cell-wise and module-wise comparisons did not appear to be related to statistical power.

Interestingly, we found an increase in the unexpectedly positive and significant correlations using both the maximum penalty score and maximum penalty ratio when a multiple myeloma dataset was used as input for scFEA and Compass with their respective imputation methods enabled. This was accompanied by a greater negative cell-wise correlation. The overall profile of stronger negative cell-wise agreement, but lower module-wise agreement was similar to our bulk data results. Thus, it appears that as cells become more homogenous in their expression profile, with potentially fewer zero values, better cell-wise similarity between Compass and scFEA becomes apparent. The increase in cell-wise consensus, at the cost of module-level consensus, was somewhat unexpected. The most likely explanation is that genetic homogeneity between cells reduces coarse trends in predicted flux across every cell, but facilitates consensus for any particular cell, possibly through the reduction of zero values. Thus, imputation measures should be approached with caution when interpreting the results of scFEA and Compass when run in concert.

### Module-wise consensus across biological processes

We were able to use our module-wise comparisons to address whether the systems broadly agreed across the results for healthy or pathological states, and for which biological processes. It has been previously speculated that metabolic perturbation could limit the usefulness of FBA methods operating under the steady-state assumption [1, 47]. Our comparison results contradicted this hypothesis. We found a high general level of module-wise agreement between the two systems for a multiple myeloma dataset, and for pancreatic cancer cell data. Relative to healthy cells, cancer cells are known to upregulate glycolysis and glutamine uptake which fuels increased nucleotide synthesis [48]. For these specific systems, we found that glycolysis and TCA, and broad sections of pyrimidine metabolism, showed greater agreement for the multiple myeloma than for a healthy mouse dataset. Thus, our current results support the use of Compass-to-scFEA consensus measures when studying single-cell tumor cell metabolism, although the systems may exhibit limited agreement on particular reactions related to nucleotide synthesis in these cells.

### Reaction-level consensus and consistency with module-level analyses

scFEA outputs flux through an entire module, where the flux for each reaction within a module can be conceptualized as contributing to the overall flux through the module [1]. Compass penalties are at the reaction level, and this calculated likelihood of a reaction is not used in additional balance calculation [5]. Thus, Compass appears to have the ability to identify individual inhibitory or potentiating reactions that could drive overall flux at steady state in a physical system. Because Compass outputs are at the reaction level, we sought to determine whether the results of scFEA and Compass showed consistency across a broad range of individual reactions related to a particular module or group of modules.

Our reaction level analysis showed a strong coherence with our representative module penalty methods. Across our single-cell datasets, and for the vast majority of individual glycolysis and TCA reactions, we found significant and negative correlations between the forward reaction penalties of Compass and the predicted fluxes of scFEA. The minority of Compass reactions that were incorrectly directed relative to an scFEA module did not appear to affect the results, potentially owing to a proportionate movement between the forward and reverse reactions, even within branched networks. For glycolysis in particular, our results suggested a stronger consensus between the two systems for the tumor dataset relative to the control, again in contrast to the idea that the steady-state assumption will not be applicable in cancers [1]. As predicted by the module-level analysis, the purine synthesis results were more poorly correlated with their corresponding Compass penalties, as were the totality of the bulk results. These results support a coherence between the systems at the reaction-level, at least for particular biological processes, and validate the use of the maximum penalty method at the module-level for single-cell data.

Our reaction-level analyses additionally uncovered a crucial insight regarding the positive correlations observed at the module-level which helped to resolve our question of how best to measure disagreement between the systems. We found that for the single-cell results, the positively correlated fluxes observed at the module-level were also observed at the reaction-level. However, at the reaction level, most of these fluxes showed universally positive correlations between the forward Compass reaction penalties and an associated scFEA predicted flux across an entire Recon subsystem. These results strongly suggest that the positive correlations observed at the module level represent a true disagreement between Compass and scFEA, and do not arise as a result of our comparison method limitations, such as unmapped reactions. Thus, this type of trend may be a strong indicator of genuine disagreement between Compass and scFEA, and the results for such reactions should be cautiously interpreted.

Given these findings, our reaction and module-level analyses suggest a complementary use of Compass and scFEA. A module-level analysis can demonstrate agreement between the two systems within a particular biological process, or spanning all available processes. Agreement within these classes could suggest whether a higher resolution analysis at the reaction level is appropriate, and where a constrained interpretation is warranted. For areas of consensus between the two systems, the reaction-level fluxes driving the overall movement at the pathway-level in biological system are suggested with a higher degree of confidence. Such a workflow might aid in identifying specific, targetable reactions that are downregulated or upregulated in a pathological state.

### Performance measures and consensus versus metabolomics testing

We tested scFEA and Compass on matched data provided by the opposing system to determine how they would perform generally, and outside of their original validation domain of either Th17 immune or pancreatic adenocarinoma cells.

Our testing showed that scFEA performed relatively well at predicting autoimmune-related metabolic alterations in mouse Th17 cells for TCA cycle and polyamine, but not glycolytic metabolism. This expands knowledge of the metabolomics-verified performance of scFEA. Further, our mouse Th17 testing suggested stronger metabolomics-validated performance for either scFEA or Compass when alternately considering glycolysis or TCA. These results suggest that the systems should ideally be run in tandem when autoimmune conditions are considered, at least for murine t-cell data. Additionally, scFEA has been used solely to predict metabolic differences in fatty acid, TCA and glycolysis between non-tumorous osteoblast precursor cells and subsequent differentiated populations associated with kidney disease, as well as a wide range of flux imbalances in tumor associated lymphocytes [49, 50]. In these studies, however, metabolomics validation appeared to be absent. Thus, expanding the biological domains in which scFEA has shown some form of metabolomics-validated predictions may be important to assessing previous and future results.

When compared to the scFEA results with the flux outlier removed, our Compass results showed a reduced ability to predict the fold-change in glycolytic and TCA cycle activity relative to metabolite data in pancreatic adenocarcinoma cells. This finding may be consistent consistent with the previous speculation that the steady-state assumption of FBA may not be amenable to the metabolic perturbation associated with tumor cells [1]. The result was somewhat unexpected given the strong negative correlations observed across both glycolysis and TCA in the multiple myeloma dataset, and the knowledge that both multiple myeloma and pancreatic adenocarcinoma cells demonstrate increased glucose and glutamine metabolism [51, 52]. One potential explanation is that Compass performs better on liquid versus solid tumor data, and that the performance decreases observed for cells originating from solid tumors could be due to gene expression alterations in a hypoxic tumor environment. [53, 54].

In this work, we did not find a statistically significant improvement of scFEA over Compass in predicting pancreatic adenocarcinoma metabolism. However, our correlation was calculated using only seven metabolites, and one substituted metabolite, in contrast to the eight used by Alghamdi et al. [1]. Further, our analysis used the entire 48 sample set of APEX1-KD knockdown data in contrast to their 27 quality-filtered cells. While our results suggest that scFEA may still be the best tool for predicting metabolism in tumor cells when clear prediction outliers are removed, it may also be that the system is sensitive to the quality of expression data originating from cells with a gene knockdown.

We assessed the level of consensus between the systems for these datasets to determine the degree to which the system predictions agreed and with the metabolomics data. For the Th17 cell data, lack of consensus between the systems, across all of the cell types and KNN settings, appeared to be reflected in the inconsistencies between the individual system predictions relative to the glycolysis and TCA metabolomics data. Thus, our consensus measures may have some ability to suggest when false positives or false negatives are present in either the Compass or scFEA results. For the pancreatic adenocarcinoma cells, significant negative correlation coefficients were observed across the glycolysis and TCA modules when the consensus tests were applied, but spanned a wide range of magnitudes. A per-module connection between the correlation coefficient and the metabolite to flux-fold prediction could not be established. However, the general agreement between the systems for the Pa03c cells, and the weak-to-moderate correlations with the metabolite data, suggests that consensus testing could reveal true positives, at least for a class of modules, indicating that the systems are consistent with the actual metabolic properties of a cell. For such a group of modules, the analysis could potentially move to the reaction-level with greater confidence. Further, although the evidence here is limited, our results suggest that module-wise consensus measures may have the ability to detect predicted flux outliers arising from scFEA, and may assist in ruling out anomalous results. This could prove useful, as single system predictions may present unusual results that might otherwise be ascribed to genuine metabolic alterations.

Given the limited availability of matched metabolomics and transcriptomics datasets, and because Compass and scFEA appeared to exhibit stronger performance for certain processes and datasets, our findings further suggest that a consensus interpretation could provide additional confidence in areas of predicted agreement and could assist in ruling out abnormal results prior to, or in absence of, any direct experimentation.

### Best practices for using scFEA and Compass with a consensus interpretation

A peripheral goal of this work was to determine whether there are additional best practices for using single-cell flux prediction tools either separately, or when consensus measures were applied. We found that the segregated FPKM datasets used in our testing produced small, but noticeable differences relative to the results of Wagner et al. It is unclear if this was the result of our using FPKM normalized counts versus the TPM normalized counts used by Wagner et al., or was caused by the data segregation itself. However, if the latter is true, it may be the case that metabolic dysregulation is not a strong enough condition to smooth data only amongst perturbed or non-perturbed cells. In terms of the consensus tests, as previously mentioned, imputation methods appeared to decrease the module-level consensus when employed by both systems. Thus, data segregation, the normalization method, and imputation should be taken into account when running these systems, particularly when their results are being compared.

### scFEA and Compass as part of a complementary workflow

Based on our findings we propose a complementary workflow using both Compass and scFEA, prior to, or in absence of metabolomics (Figure S2). Under such a workflow, both systems are run on a common single-cell dataset, and the module-level comparison tests described in this paper are performed. The module-level consensus results can be used directly to determine areas of agreement, and reaction-level consensus tests can be performed for these modules, if required. This subset of high consensus module-level or reaction-level flux predictions can undergo downstream analysis using either the scFEA or Compass predictions.

### Limitations

Several limitations were present in this study. Our automated method of clustering and selecting Compass penalties to represent a module appeared to be successful in practice. However, the mapping of Recon reaction identifiers to most scFEA modules did not produce the same number of reactions as in the original module. In cases where either a KEGG id or EC number mapped to multiple reactions or reaction variants, these could not readily be filtered from the results. Further, an exhaustive analysis of the mapped reactions was not performed.

As this work focused on the methods of comparing the outputs of the two systems, the single-cell RNA-seq datasets used were limited. Alternative matched metabolomics and single-cell RNA-seq datasets were not readily available. Similarly, an exhaustive analysis comparing the Compass predicted reaction penalties for each Recon2 subsystem to its scFEA analog was not undertaken. Thus, it is not certain how our results would generalize to further datasets and reaction subsystems.

Owing to the lack of matched metabolomics and single-cell RNA sequencing data available, mouse data was used in this study during consensus testing. While scFEA provides explicit support for mouse data, a published listing of the reactions for each of the mouse modules does not appear to be available. We were able to work around this limitation by using the human module information from scFEA, under the assumption that the reactions should be similar for both species. Consensus measures based on the 168 human module metadata were applied to the 168 module scFEA mouse results, but it is not explicitly known if these modules are analogous. Further, Compass uses the Recon2 human metabolic reconstruction, and although the effects are assumed to be minor, it is not known to what degree this inherent limitation affected our mouse results.

Lastly, Compass penalties were selected for each module based on a maximum for non-identical but related reactions. These reactions may have dissimilar optima as determined by the first stage of Compass algorithm. Therefore, although our method appeared to be successful, it is not precisely known how the reaction optima calculated by this initial FBA step can influence the penalty versus the transcriptome-inclusive second step.

## Conclusing remarks and future work

Here, we demonstrated that the similarities between outputs of Compass and scFEA can be measured, and that these systems can achieve a substantial consensus for at least certain types of single-cell data. The complementary nature of the two systems can allow movement to the reaction-level if agreement is observed at the broader module level. These systems appear to show a stronger agreement when applied to RNA sequencing data from metabolically disturbed cells. Optimal use of these systems, particularly together, may require extra data preparation steps and runs with different system settings. Homogenizing the expression profiles of cells may detrimentally affect the ability of this methods described in this work to detect consensus.

In this work, it was not possible to extensively test large combinations of datasets and software settings. Future work might determine the level of consensus between Compass and scFEA for datasets arising from a range of species, pathological and healthy states and sequencing technologies. Further investigating best practices, such as imputation settings, dataset segregation, and standardization of library and linear solver versions should also be implemented if this method is to be broadly adopted.

To improve the consensus, enhanced mapping methods could be used to remove unwanted compartmental variants and reactions from the module-to-reaction key. Manual verification could be used to ensure that each group of reactions was accurately mapped. To better compare the penalties of disparate and remotely related reactions, Compass penalty scores could be normalized for their optima so that difficult to maintain low optima reactions can be differentiated from easily maintained high optima reactions. In addition, determining the effect of substrate substitutions on a wide range Compass reaction penalties could also suggest whether they should be filtered from the reaction collections.

Beyond optimizing the use of current tools, our paper suggests a path towards improving these single-cell methods. Previously, prediction of metabolism by models based on transcriptomic data has shown highly conflicting results. Using data from the Cancer Cell Line Encyclopedia to train a co-occurrence network, Caviccioli et al. demonstrated that metabolites could be predicted with varying levels of success [55]. However, Li et al. reported negative results when using a lasso regression trained on the same dataset [56]. Notably, however, Li et al. showed marginal increases in predictive performance when their model was given upstream metabolites in addition to transcriptomics data, suggesting that that the order and position of metabolites is important to prediction [56].

The consensus between Compass and scFEA observed in this work, as well as their level of performance here and in their respective studies, further suggests that combining metabolic network information with initially weak pathway flux predictions may enhance flux estimations across the reactions of an entire metabolic model. Indeed, earlier literature on flux prediction supports the requirement, and suggests that combining a metabolic network with transcriptomics could result in an over 10-fold increase in accuracy [57].

The co-occurrence network approach of Caviccioli et al., as well an investigation of the relationship between gene expression and metabolite quantities, suggests that non-pathway genes can potentially have a greater influence over metabolite levels than pathway genes alone [58]. We were able to verify this with our own experiments. Thus, to improve upon existing methods, two conditions are suggested. First, there should be convincing evidence of gene-metabolite correlations, even if the predictive genes are not coding for pathway enzymes directly related to the metabolite. Second, the ability of a chosen gene-set to predict metabolite quantity should be closely surveyed as metabolic network information is incorporated into the model.

## Supporting information

**Tables S1-S4**. Cell and module similarity measures for two methods of comparing Compass penalties to the predicted fluxes of scFEA for three RNA-seq datasets.

**Table S1.** Single-cell multiple myeloma gene expression dataset (GSE110499).

| Cell |  |  |  |  |
| --- | --- | --- | --- | --- |
| Aggregating function | Penalty Ratio |  | Positive Penalty |  |
|  | r | P<sub>comb</sub> | r | P<sub>comb</sub> |
| Max | -0.054 | 0.79 | -0.066 | 0.091 |
| Min | -0.0018 | 1 | -0.0033 | 1 |
| Sum | -0.081 | 0.0012 | -0.068 | 1 |
| Mean | -0.18 | 0.30 | -0.070 | 0.82 |
| Module |  |  |  |  |
| Aggregating function | Penalty Ratio |  |  |  |
| | p < 0.05 | NA | Positive $\rho$ | $\frac{\text{Positive } \rho}{p < 0.05}$ |
| Max | 103 | 7 | 11 | 0.11 |
| Min | 44 | 70 | 4 | 0.085 |
| Sum | 103 | 7 | 11 | 0.11 |
| Mean | 103 | 7 | 11 | 0.11 |
| Aggregating function | Positive Penalty |  |  |  |
| | p < 0.05 | NA | Positive $\rho$ | $\frac{\text{Positive } \rho}{p < 0.05}$ |
| Max | 106 | 5 | 9 | 0.085 |
| Min | 54 | 67 | 5 | 0.093 |
| Sum | 107 | 5 | 9 | 0.084 |
| Mean | 107 | 5 | 9 | 0.084 |

**Table S2.** Single-cell fetal mouse lung dataset (GSE122330).

| Cell |  |  |  |  |
| --- | --- | --- | --- | --- |
| Aggregating function | Penalty Ratio |  | Positive Penalty |  |
|  | r | P<sub>comb</sub> | r | P<sub>comb</sub> |
| Max | -0.059 | 0.92 | -0.084 | 0.0053 |
| Min | -0.051 | 1 | -0.048 | 1 |
| Sum | -0.067 | 0.30 | -0.064 | 0.30 |
| Mean | -0.077 | 0.20 | -0.074 | 0.27 |
| Module |  |  |  |  |
| Aggregating function | Penalty Ratio |  |  |  |
| | p < 0.05 | NA | Positive $\rho$ | $\frac{\text{Positive } \rho}{p < 0.05}$ |
| Max | 77 | 14 | 3 | 0.039 |
| Min | 26 | 77 | 4 | 0.15 |
| Sum | 82 | 14 | 4 | 0.049 |
| Mean | 82 | 14 | 4 | 0.049 |
| Aggregating function | Positive Penalty |  |  |  |
| | p < 0.05 | NA | Positive $\rho$ | $\frac{\text{Positive } \rho}{p < 0.05}$ |
| Max | 82 | 12 | 3 | 0.037 |
| Min | 39 | 73 | 3 | 0.077 |
| Sum | 90 | 12 | 5 | 0.056 |
| Mean | 90 | 12 | 5 | 0.056 |

**Table S3.** Single-cell multiple myeloma dataset (GSE110499) with imputation and smoothing.

| Cell |  |  |  |  |
| --- | --- | --- | --- | --- |
| Aggregating function | Penalty Ratio |  | Positive Penalty |  |
|  | r | P<sub>comb</sub> | r | P<sub>comb</sub> |
| Max | -0.010 | $9.13 \times 10^{-12}$ | -0.011 | $1.15 \times 10^{-17}$ |
| Min | 0.036 | 1 | 0.035 | 1 |
| Sum | -0.11 | $4.75 \times 10^{-19}$ | -0.11 | $1.30 \times 10^{-16}$ |
| Mean | -0.00038 | 1 | -0.007 | 1 |
| Module |  |  |  |  |
| Aggregating function | Penalty Ratio |  |  |  |
| | p < 0.05 | NA | Positive $\rho$ | $\frac{\text{Positive } \rho}{p < 0.05}$ |
| Max | 106 | 11 | 30 | 0.28 |
| Min | 47 | 74 | 12 | 0.25 |
| Sum | 108 | 11 | 30 | 0.28 |
| Mean | 108 | 11 | 30 | 0.28 |
| Aggregating function | Positive Penalty |  |  |  |
| | p < 0.05 | NA | Positive $\rho$ | $\frac{\text{Positive } \rho}{p < 0.05}$ |
| Max | 109 | 9 | 28 | 0.26 |
| Min | 55 | 70 | 9 | 0.16 |
| Sum | 110 | 9 | 28 | 0.25 |
| Mean | 110 | 9 | 28 | 0.25 |

**Table S4.** Bulk Ovarian Clear-cell Carcinoma (GSE123426).

| Cell |  |  |  |  |
| --- | --- | --- | --- | --- |
| Aggregating function | Penalty Ratio |  | Positive Penalty |  |
|  | r | P<sub>comb</sub> | r | P<sub>comb</sub> |
| Max | -0.25 | $1.29 \times 10^{-17}$ | -0.21 | $2.0 \times 10^{-11}$ |
| Min | 0.14 | $2.51 \times 10^{-4}$ | 0.16 | $5.1 \times 10^{-6}$ |
| Sum | -0.26 | $4.75 \times 10^{-19}$ | -0.25 | $1.30 \times 10^{-16}$ |
| Mean | -0.0054 | 0.90 | -0.015 | 1 |
| Module |  |  |  |  |
| Aggregating function | Penalty Ratio |  |  |  |
| | p < 0.05 | NA | Positive $\rho$ | $\frac{\text{Positive } \rho}{p < 0.05}$ |
| Max | 27 | 11 | 8 | 0.30 |
| Min | 9 | 75 | 0 | 0.00 |
| Sum | 24 | 11 | 4 | 0.17 |
| Mean | 24 | 11 | 4 | 0.17 |
| Aggregating function | Positive Penalty |  |  |  |
| | p < 0.05 | NA | Positive $\rho$ | $\frac{p < 0.05}{\text{Positive } \rho}$ |
| Max | 28 | 9 | 6 | 0.21 |
| Min | 12 | 70 | 1 | 0.083 |
| Sum | 23 | 9 | 2 | 0.087 |
| Mean | 23 | 9 | 2 | 0.087 |

**Fig S1.**
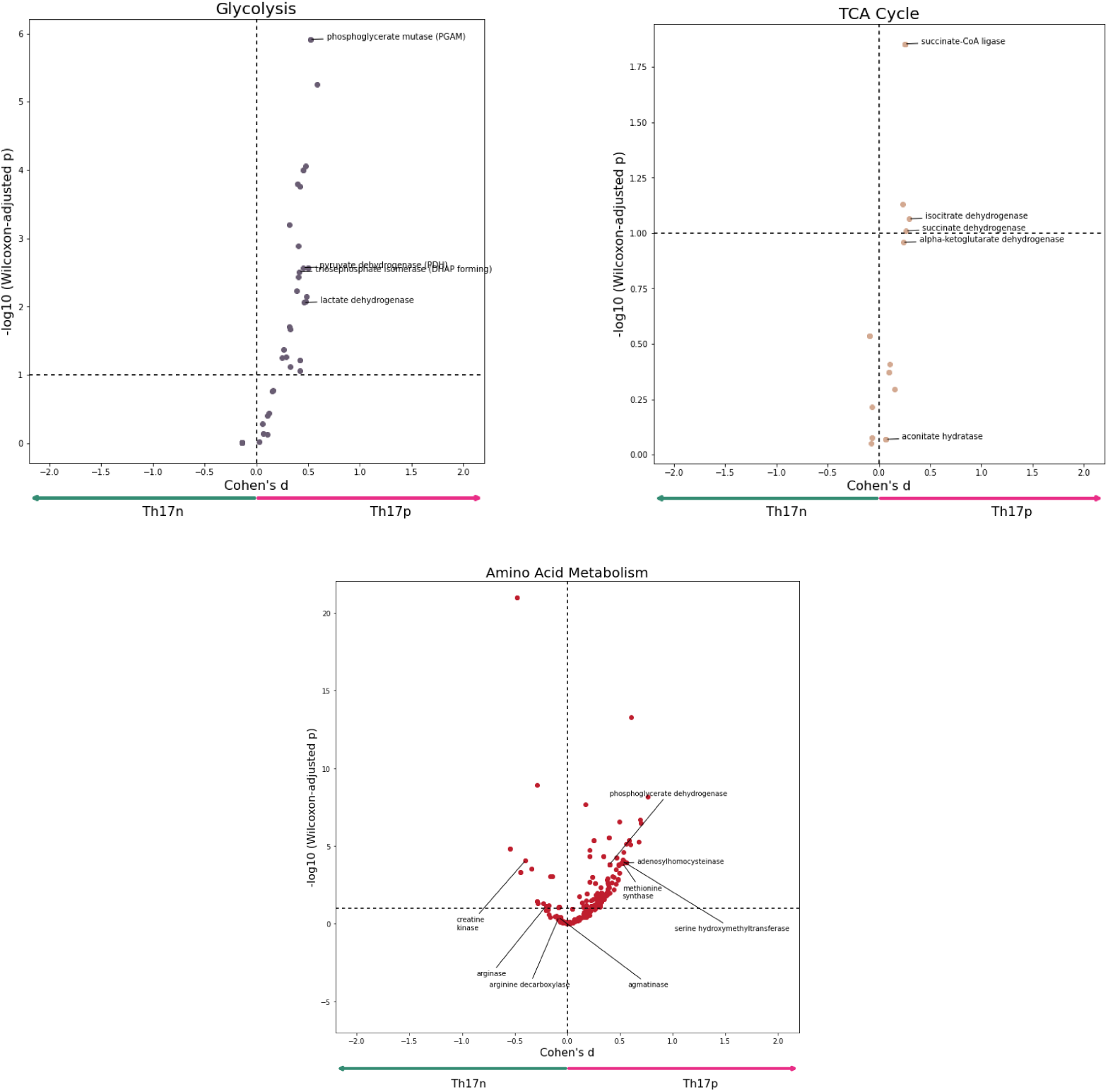
The glycolysis, TCA, and polyamine results from adapting the post-processing code of Wagner et al. to use separate pathological and normal Th17 results. Each point represents single directed reaction. The effect size between the two cell types is shown by the Cohen’s distance on the x-axis. The y-axis is the negative log of the adjusted p-value for a Wilcoxon test between identical reactions in each cell type.

**Fig S2.**
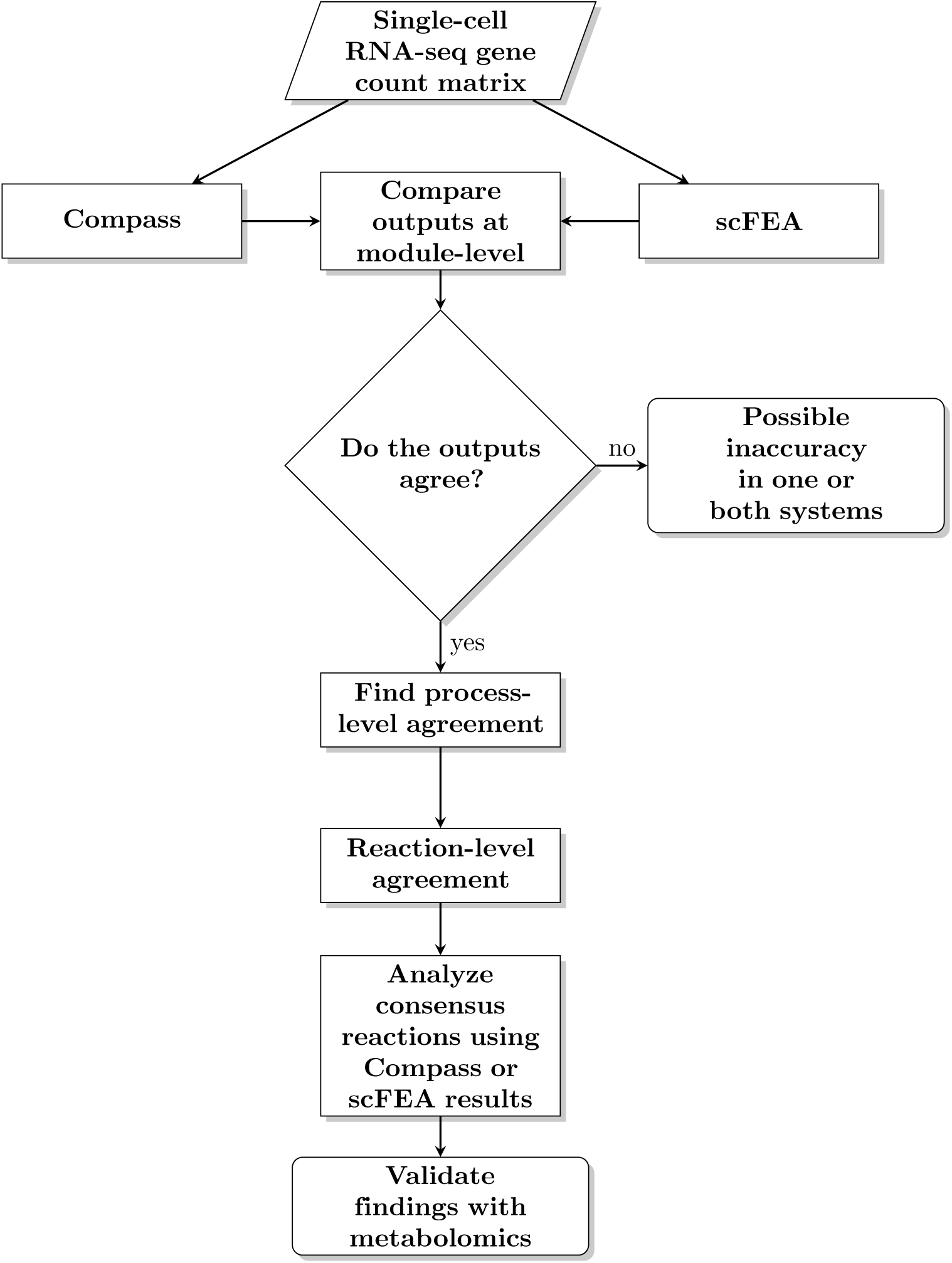
A flowchart of a potential single-cell metabolic prediction workflow using scFEA, Compass and our module-level consensus test. The outputs of the systems are globally compared and reduced down to a set of consensus reactions. Among these consensus reactions, a researcher can select relevant reactions, and analyze the corresponding scFEA and Compass results. Metabolomics validation could follow.

## S1 Appendix. A simplified representation of the first two Compass algorithms

The steps used by Compass to convert normalized gene count data to reaction penalties are shown in the preparatory steps. The transcript-agnostic flux-balance analysis step is used to determine the optimum flux for each reaction, subject to a system-wide steady-state condition. In the transcript-inclusive step, the probability of a reaction is calculated by incorporating the reaction penalties scores from the data preparation stage into a cell-wise objective function. The values chosen here are not based on actual transcriptomics data, or intermediates from Compass, but were selected for convenience to give an intuition into the first two Compass algorithms.

### Preparatory Steps

1. Single-cell gene expression matrix input:

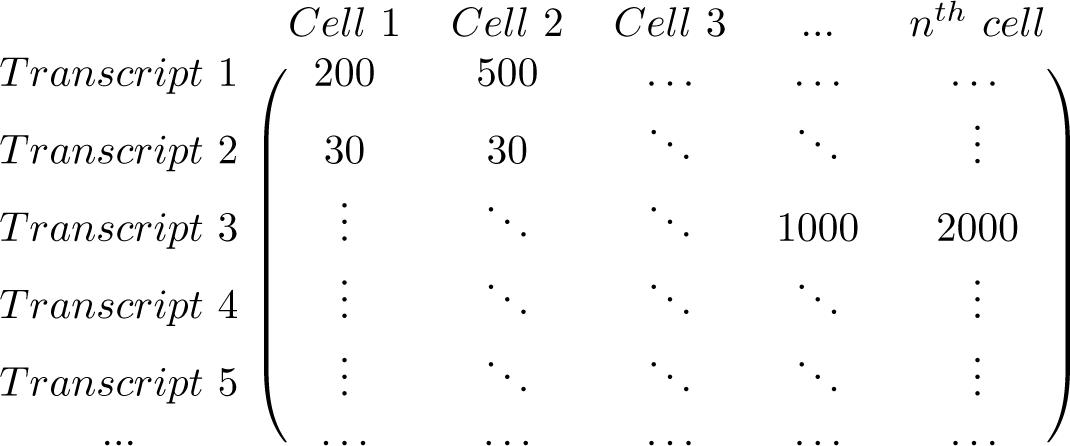
2. Map gene expression to reactions to create reaction expression matrix R(G). Multiple genes can be assigned to single reactions in a boolean- wise fashion, as in the case of multi-subunit enzymes.

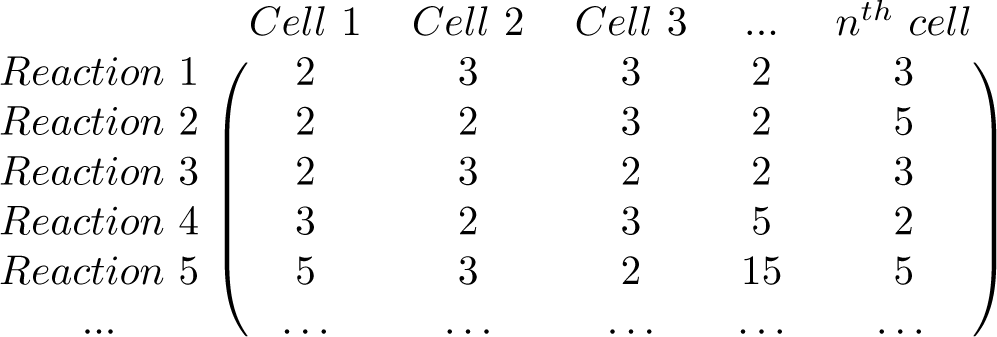
3. Take the inverse of each element to obtain reaction penalty matrix P(G):

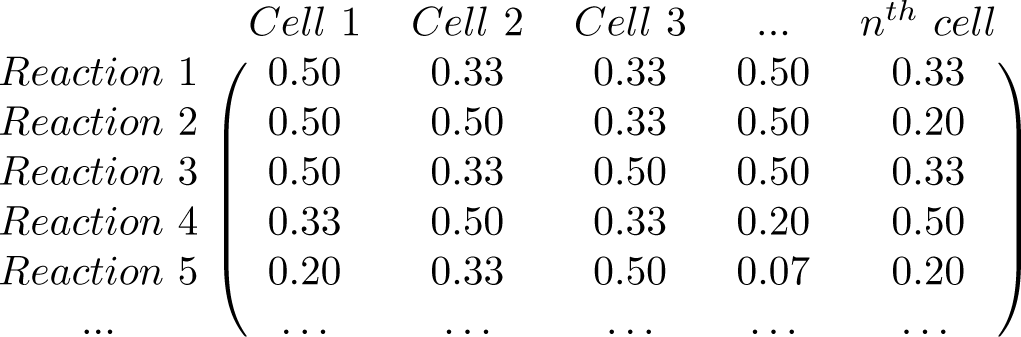

### Standard Flux-Balance Step (No Gene Inputs)

1. Set up standard flux-balance equations using stochiometric matrix with metabolites (*A − E*) and reactions (*R*_1_ *− R_n_*).

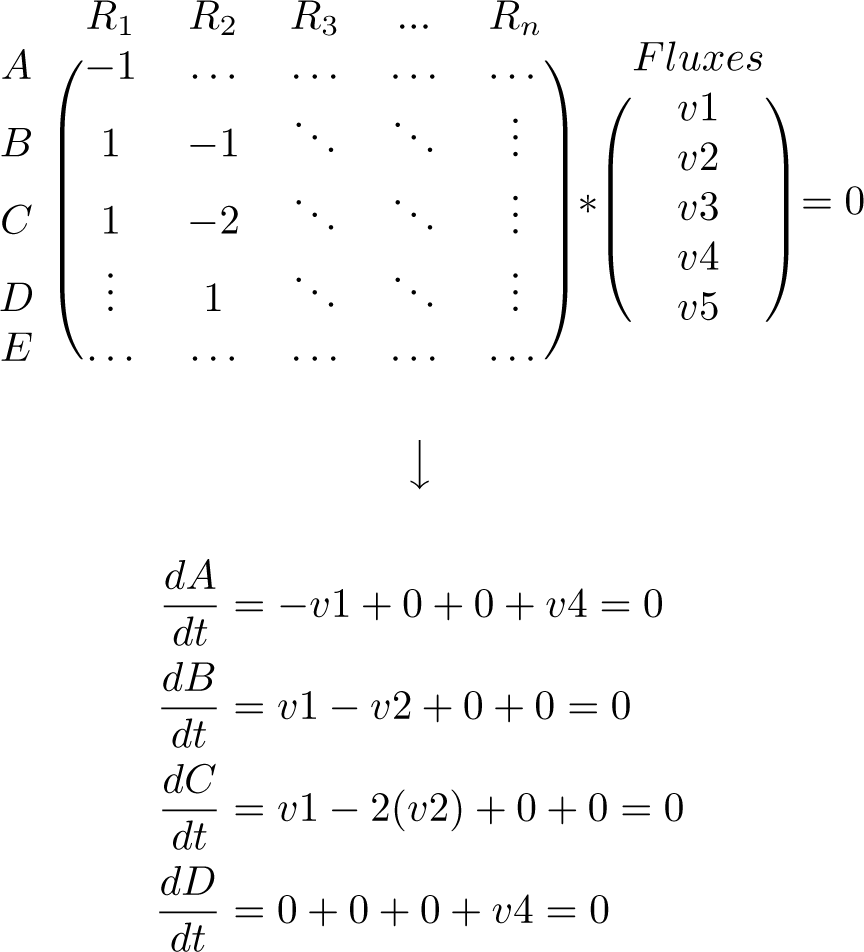
2. Calculate the optimum flux for each reaction (v*_opt_*) by iterating through each reaction, setting a single as the objective, and maximizing it.

### Transcript-inclusive flux-balance analysis step

1. Create a cell-wise objective function by multiplying each flux variable by the reaction penalties in matrix P(G) for one cell:

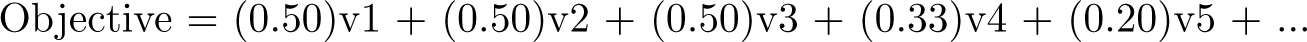
3. Select first reaction.

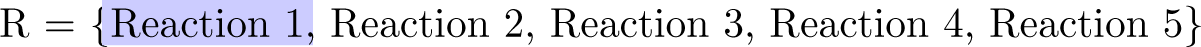
4. Hold flux associated with reaction within 95% of its optimum from the standard FBA step and minimize the objective function.

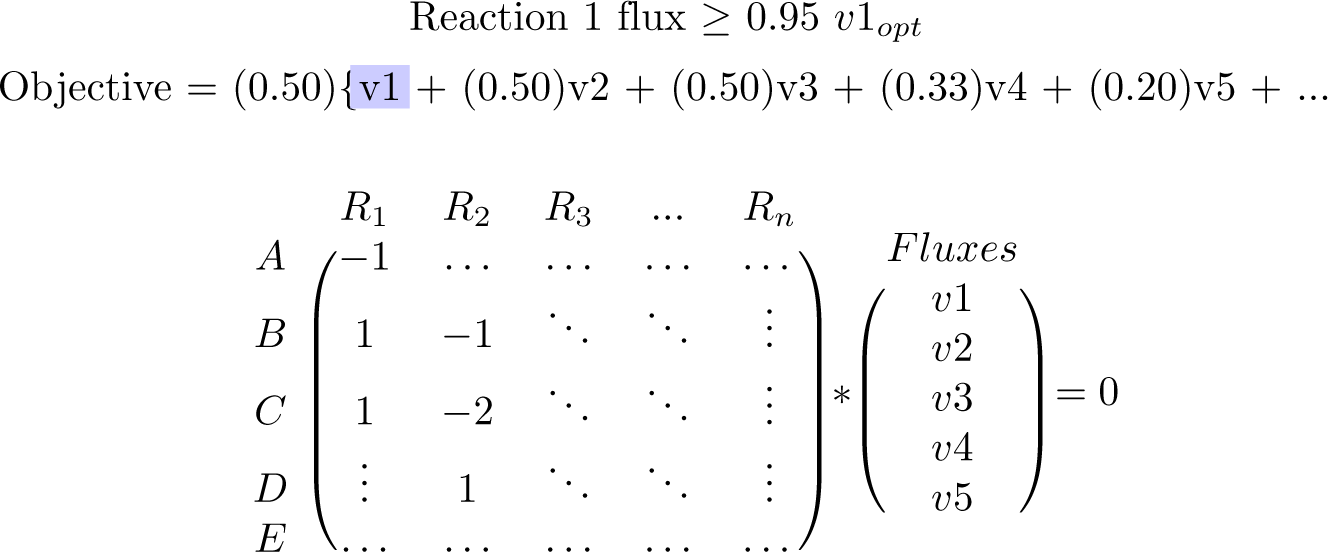
5. The solution is the minimum global flux, subject to transcript limitations, required to maintain the optimum for the selected reaction (lower is more likely). Iterate through the reactions, repeating steps 1-4.

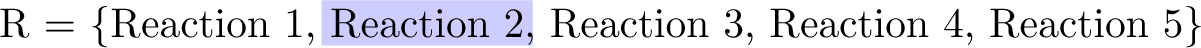

## S2 Appendix. A simplified example of the preparatory steps in flux estimation taken by scFEA

A simple four-step linear reaction system is used to illustrate how the system sets up its balance conditions for use as metabolite-specific factors. Metabolite-specific rows are used to determine the flux imbalance, the sum of all fluxes producing a metabolite versus the sum of all fluxes consuming the metabolite.

### scFEA Preparatory Steps

1. A representation of a simple linear reaction chain where metabolites A-C are the products or initial reactants of the reactions within modules M1-M4.

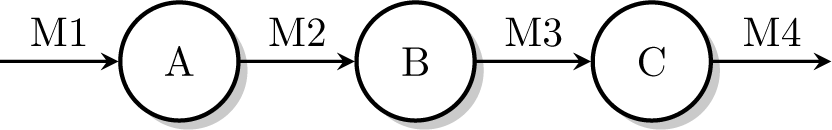
2. The modularized stochiometric matrix for the above reaction chain.

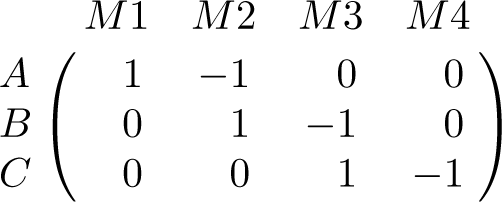
3. The matrix with flux variables multiplied in and separated into metabolite-specific rows where each row will be used as a factor.

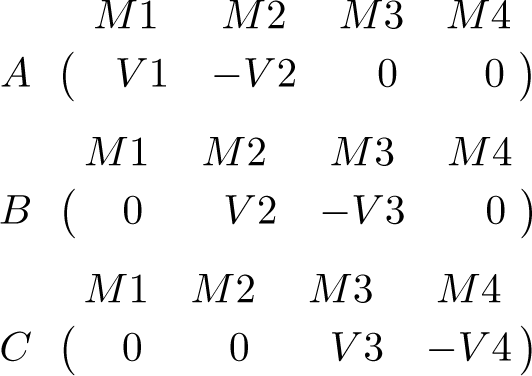
4. Each flux *V* 1-*V* 4 is estimated by a neural subnetwork that takes as inputs the genes associated with the corresponding module. During training, the loss function will minimize (in part) the squared sum of the flux imbalances associated with each metabolite across the entire factor graph.

